# Human-in-the-loop optimization of visual prosthetic stimulation

**DOI:** 10.1101/2021.11.24.469867

**Authors:** Tristan Fauvel, Matthew Chalk

**Affiliations:** Institut de la Vision Sorbonne Université, INSERM, CNRS 17 rue Moreau, F-75012, Paris, France

## Abstract

Retinal prostheses are a promising strategy to restore sight to patients with retinal degenerative diseases. These devices compensate for the loss of photoreceptors by electrically stimulating neurons in the retina. Currently, the visual function that can be recovered with such devices is very limited. This is due, in part, to current spread, unintended axonal activation, and the limited resolution of existing devices. Here we show, using a recent model of prosthetic vision, that optimizing how visual stimuli are encoded by the device can help overcome some of these limitations, leading to dramatic improvements in visual perception. We propose a strategy to do this in practice, using patients’ feedback in a visual task. The main challenge of our approach comes from the fact that, typically, one only has access to a limited number of noisy responses from patients. We propose two ways to deal with this: first, we use a model of prosthetic vision to constrain and simplify the optimization; second, we use preferential Bayesian optimization to efficiently learn the encoder using minimal trials. To test our approach, we presented healthy subjects with visual stimuli generated by a recent model of prosthetic vision, to replicate the perceptual experience of patients fitted with an implant. Our optimization procedure led to significant and robust improvements in perceived image quality, that transferred to increased performance in other tasks. Importantly, our strategy is agnostic to the type of prosthesis and thus could readily be implemented in existing implants.

## 1 Introduction

Retinal prosthetics aim at restoring vision by electrically stimulating neurons in the retina (Lorach et al., 2013). In existing devices, the patient wears a glasses-mounted camera that captures the visual scene. This visual input is then transformed by a small pocket computer into patterns of electrical pulses. Despite recent progress (Palanker et al., 2020), the visual resolution that can be achieved with such implants is very low: below the threshold for legal blindness (Stingl et al., 2017; Palanker et al., 2020). This is due, in particular, to the limited spatial resolution of stimulating currents, the activation of passing axons, and the unspecific stimulation of cells of different types (Weitz et al., 2015; Grosberg et al., 2017; Golden et al., 2019; Beyeler, 2018; Beyeler et al., 2019).

To improve visual performance, recent research has sought to optimize how the prosthetic device converts visual stimuli into patterns of electrical stimulation. One approach has been to try to replicate the inputs that retina ganglion cells (RGCs) receive in normal conditions (Nirenberg and Pandarinath, 2012; Shah et al., 2019). However, this requires a model that can well predict RGC responses to natural stimuli, which is currently lacking (McIntosh et al., 2016). Recent work has sought to develop stimulation protocols that can selectively activate different cell types, in a manner akin to natural stimulation (Twyford et al., 2014; Guo et al., 2018; Lee and Im, 2019), but the limited resolution and signal-to-noise ratio achievable with existing devices make it difficult to match the activity of RGCs under natural conditions. Moreover, these methods often depend on being able to simultaneously record and stimulate RGC responses and thus would require significant technological advancements to be used on patients (Shah and Chichilnisky, 2020; Rincón Montes et al., 2020).

Another approach is to use feedback from patients to optimize how visual inputs are encoded by the device (Eckmiller et al., 1999; Becker et al., 1999; Eckmiller et al., 2005, 2007). A potential advantage of this approach is that the encoder can adapt to deal with differences in the visual perception reported by different patients (Beyeler et al., 2019). On the other hand, it is limited by the fact that psychophysical data are typically noisy and slow to collect (Barnes et al., 2016).

Here we propose a new strategy to optimize how visual stimuli are encoded, based on patients’ responses in a visual task. The main challenge we faced was to optimize a high-dimensional encoder based on limited, noisy, responses from patients. We used two complementary approaches to address this. First, we used a biologically constrained model, describing how patterns of electrical stimulation are converted to a visual percept, to reduce the number of parameters of the encoder and simplify the optimization (Nanduri et al., 2012; Beyeler et al., 2019). Second, we developed a novel preferential Bayesian optimization algorithm to efficiently optimize the encoder using a limited number of patients’ responses.

We considered a simple visual task, in which patients are presented with a single letter, and have to compare which of two different encoders generated a less distorted visual percept. Their responses on each trial were used to optimize the encoder, so as to progressively increase the quality of the resulting visual percept. To demonstrate how our approach could work in practice, we presented healthy subjects with visual stimuli generated using a computer model, designed to mimic the perceptual experience of patients fitted with a prosthetic implant. Our approach led to a significant and robust improvement in perceived image quality, that transferred to improvements in other tasks. As our approach requires no new hardware, it should be relatively easy to implement with existing technologies.

## 2 Methods

We consider a static image, denoted by an *n*_*s*_ × 1 vector, *s* (where *n*_*s*_ is the number of recorded image pixels). This image is transformed by the prosthetic device into a pattern of electrical stimulation, denoted by an *n*_*e*_ × 1 vector *a* (where *a*_*i*_ is the current amplitude of the *i*^*th*^ electrode and *n*_*e*_ is the total number of electrodes). We define a retinal prosthetic ‘encoder’ as the mapping from the stimulus *s* to amplitudes *a*. For simplicity, we assumed a linear encoder, *a* = ***W****s*, where ***W*** is an *n*_*e*_ × *n*_*s*_ matrix of encoding weights. Rows of ***W*** can be interpreted as the‘receptive field’ of each electrode, i.e., how much each image pixel contributes in stimulating a given electrode.

Our goal is to find a set of weights, ***W***, that maximizes the perceptual quality experienced by the patient for any input, *s*. We consider that the only available data are subjects’ perceptual reports, and we aim at learning the best encoder by measuring the subject’s responses in visual tasks with different encoders. For simplicity, we consider a static model, where the electrical pulse shapes and frequency are kept fixed, and only the pulse amplitudes vary, but our approach could generalize to include these variables.

### 2.1 Experiments on normally sighted subjects

To develop our approach, we experimented on healthy subjects using the prosthetic vision simulator *pulse2percept* (**Fig 1A**; Beyeler et al. (2017a)). This model simulates the visual percepts experienced by patients fitted w ith a prosthetic device. In particular, it describes the effects of current diffusion and activation of passing axons on the perceived phosphenes. It was fitted and validated using data from users of the epiretinal implants Argus I and Argus II (Second 79 Sight Medical Products, Sylmar, CA) (Horsager et al., 2009, 2010, 2011; Nanduri et al., 2012; Beyeler et al., 2019). Here, we focused on Argus II, a 60-electrode epiretinal device that has been implanted in approximately 300 individuals since its commercial approval in the EU in 2011, the US in 2013, and Canada in 2015 (Ahuja et al., 2011; Humayun et al., 2012; Rizzo, 2011; da Cruz et al., 2016).

**Figure 1:**
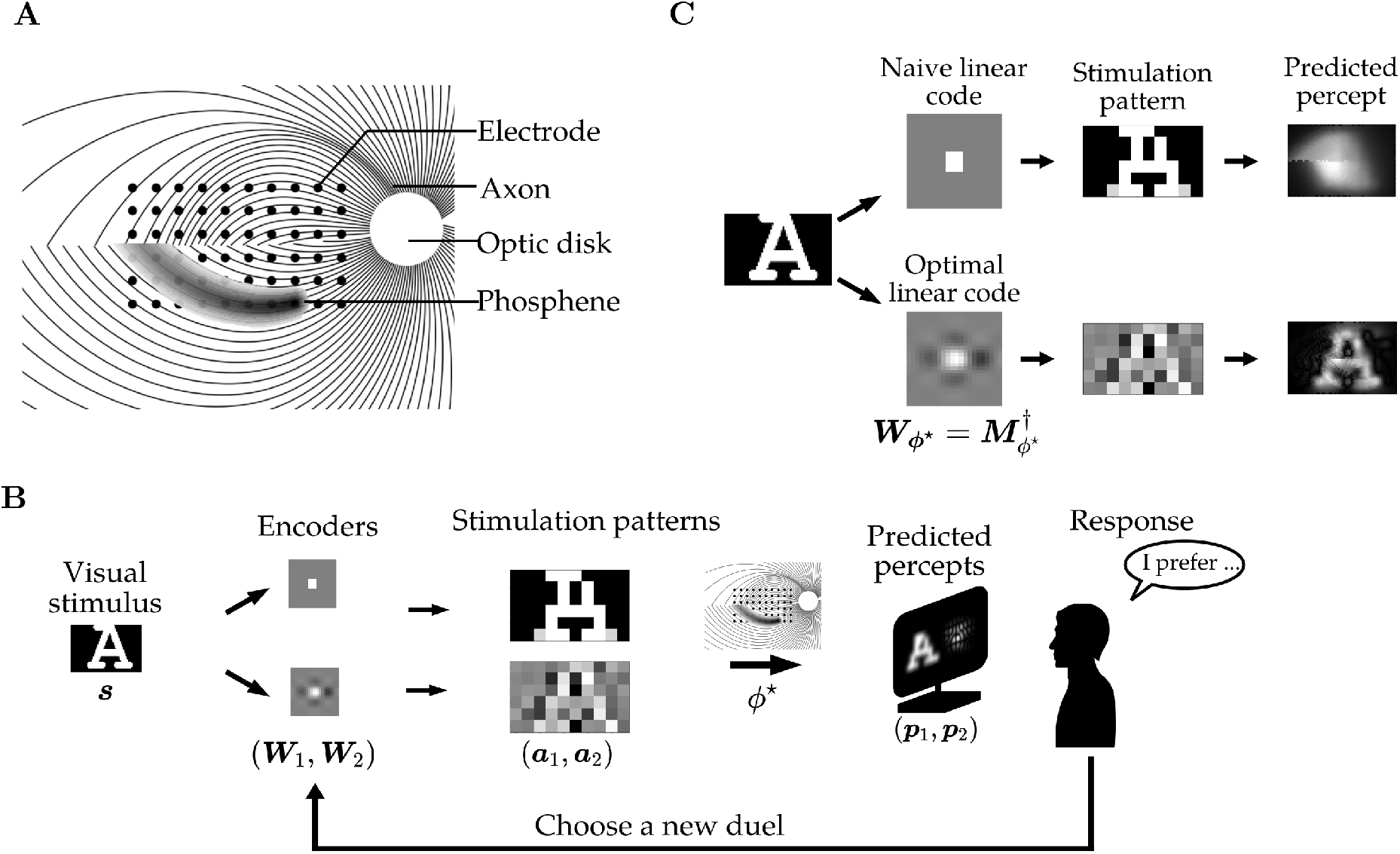
Framework. (**A**) Schematic of pulse2percept model. An electrode array (black dots) stimulates the retina. In addition to stimulating nearby neurons, electrodes stimulate passing axon fibers that project towards the optic disk (curved lines). Consequently, the phosphene induced by a single electrode has a ‘comet shape’, following the patch of nearby axons (shaded black). Eight parameters (ϕ), corresponding to the position of the implant, the electrical properties of the retina, and the axon map, determine the percept that results from a given stimulation pattern. (**B**) Preference task. A stimulus s is converted via two different encoders (**W**_1_, **W**_2_) into two different stimulation patterns (a_1_, a_2_). Predicted percepts, generated using the pulse2percept model with parameters ϕ*, are displayed to subjects onscreen. Subjects are asked which of the two percepts they find less d istorted. Their responses are used to decide which encoders to present on the next trial, to incrementally improve the simulated percepts. (**C**) Comparison of a naive encoder, where the image is simply down-sampled, to an optimal encoder (optimized for a given, known, perceptual model, ϕ*). The ‘receptive field’ ( RF) of a single electrode ( how much each image pixel contributes in stimulating this electrode) is shown for both encoders. Optimizing the encoder results in center-surround RFs, where electrodes are suppressed by visual illumination close to the center of their RF. This allows compensating for distortion due to current spread and axon activation (panel A), resulting in a dramatic improvement in the predicted percept.

The *pulse2percept* model assumes that input electrode currents combine linearly to produce a spatial intensity profile *I*(*x, y*), where (*x, y*) are spatial coordinates on the retina (Beyeler et al., 2019). This intensity profile is transformed linearly to take into account axonal activation, before going through a static nonlinearity, to obtain subjects’ predicted percepts. Thus, the static version of *pulse2percept* is essentially a linear-non-linear model.

Let *ϕ* denote the parameters describing the *pulse*2*percept* model for a specific subject. The visual percept predicted by the model, *p* (an *n*_*p*_ × 1 vector, where *n*_*p*_ is the number of pixels in the percept), is given by:

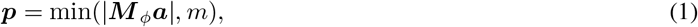

where *m* limits the maximum brightness of the percept (we set *m* = 1), and *M*_*ϕ*_ is an *n*_*p*_ × *n*_*e*_ matrix whose elements depend on various parameters of the *pulse2percept* model, *ϕ*. Rows of *M*_*ϕ*_ correspond to electrodes’ ‘projective fields’: the linear contribution of a single electrode to the visual percept.

Eight patient-specific parameters *ϕ* were used to parameterize the encoder: the current diffusion coefficient, the axon sensitivity, the implant location and orientation, and two parameters describing the trajectory of axons (Jansonius et al., 2009). A last parameter was used to scale the current intensity at each electrode. Based on these parameters, *pulse*2*percept* predicts the projective field of each electrode in *M*_*ϕ*_.

Since this formulation of the *pulse*2*percept* model is not apparent in the original publication of Beyeler et al. (2017a), we compared the percepts predicted by their code to those predicted with our simpler linear-nonlinear formulation. We generated 10^4^ random amplitude patterns and computed the difference in the predictions of the two versions of the model, keeping all parameters of the *pulse*2*percept* model constant, at their default values. The root-mean-squared error was 2.8 × 10^−5^, implying that the two versions indeed correspond to the same model.

In previous work, patients fitted with the Argus I I device reported a range of different visual percepts for identical stimulation currents, likely due to individual variations in the position and electrical properties of their implant (Beyeler et al., 2019). To replicate this, we varied the parameters used to generate the simulated visual percept for each subject (parameters were sampled randomly in the ranges shown in table 1). Note, however, that the parameters determining the axon shape descriptors were kept constant, as they were later modified to assess the robustness of our approach to a wrong parameterization (section 3.5).

**Table 1:** Search space. ρ is the current diffusion coefficient, λ is the axons sensitivity to current. ρ and λ ranges are taken from Beyeler et al. (2019). x and y are the implant coordinates with respect to the fovea. θ is the implant orientation with respect to the horizontal axis. β_−_ and β_+_ are axons shape descriptors, their values are taken from Jansonius et al. (2009). The amplitude parameter scales the percept brightness.

| | $\rho$ ($\mu\text{m}$) | $\lambda$ ($\mu\text{m}$) | $x$ ($\mu\text{m}$) | $y$ ($\mu\text{m}$) | $\theta$ | $\beta_-$ | $\beta_+$ | Amplitude (arb. unit) |
| --- | --- | --- | --- | --- | --- | --- | --- | --- |
| Lower bound | 137 | 358 | -500 | -500 | -15° | 0.1 | -2.5 | 0 |
| Upper bound | 415 | 1510 | 500 | 500 | 15° | 1.3 | -1.3 | 300 |

### 2.2 Encoder parameterization

We sought a low-dimensional parameterization of the encoder, ***W*** , that could be feasibly optimized using a limited number of subjects’ responses. To do this, we assumed that subjects’ percepts were generated by the *pulse2percept* model, with unknown parameters, *ϕ*. We reasoned that if these parameters are known for any given subject, then it should be possible to work out the encoder, ***W***_*ϕ*_, that results in an optimal visual percept. As a result, our problem is reduced to finding the 8 parameters of the *pulse2percept* model, *ϕ*, rather than the *n*_*e*_ × *n*_*s*_ of the encoder, ***W***.

Given *M*_*ϕ*_, we can compute the linear encoder ***W***_*ϕ*_ that minimizes the squared distance between the image *s* and the predicted percept *p*, while assuming that projective fields combine linearly (i.e. we ignore the modulus in Eqn 1):

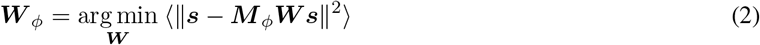

It is straightforward to show that the optimal linear encoder ***W***_*ϕ*_ is simply the pseudo-inverse of 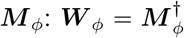. That is, the encoder ***W***_*ϕ*_ is entirely determined by the eight parameters *ϕ*, and not the stimulus, *s*.

Note that ideally, we should replace *M*_*ϕ*_***W****s* in the previous equation with min(|*M*_*ϕ*_***W****s*|, *m*). However, this would result in a more complex numerical optimization, and slow down our algorithm. In practice, numerical simulations showed that there was little difference in the percepts obtained using this approximation (Supp. Fig S1).

### 2.3 Preference-based optimization

On each trial, subjects were simultaneously presented with two different simulated percepts, generated using the *pulse2percept* model (**Fig 1B**). The percepts corresponded to the same ‘letter’ stimulus, but with two different encoders (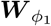 and 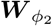). Subjects were asked to report which stimulus, corresponding to a given target letter, was the least distorted (the target letter was presented simultaneously onscreen; with real patients, we could inform subjects of the target letter via verbal instruction). The exact instructions given to participants are shown in supplementary 5. We used subjects’ reported preferences to optimize the encoder model parameter, *ϕ*.

We used a preferential Bayesian optimization (PBO) algorithm to optimize the encoder based on subjects’ reported preferences (Brochu et al., 2010; Fauvel and Chalk, 2021a). We assumed that a function *g*(*ϕ*), determines how much subjects’ ‘prefer’ a given set of parameters, *ϕ*. The probability that subjects prefer *ϕ*_1_ to *ϕ*_2_ (returning a response *ϕ*_1_ ≻ *ϕ*_2_) is assumed to depend on this preference function as follows:

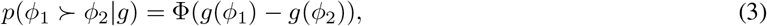

where Φ is the cumulative normal distribution function. Thus, the larger the value of *g*(*ϕ*_1_) relative to *g*(*ϕ*_2_), the more likely *ϕ*_1_ is to be preferred over *ϕ*_2_.

We developed a novel PBO algorithm to efficiently optimize *g*(*ϕ*) based on subjects’ reported preferences, called ‘Maximally Uncertain Challenge’ (Fauvel and Chalk, 2021a). As the details are described in Fauvel and Chalk (2021a), we provide only a brief overview here. As is standard in PBO, we assume a Gaussian process prior over *g*, and use an approximate inference algorithm (expectation propagation; Minka (2001); Seeger (2002)) to infer the posterior probability distribution over *g* given subjects’ reported preferences, 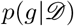. We then use this to select new parameters (*ϕ*_1_ and *ϕ*_2_) for the next trial. First, we select a ‘champion’, which maximizes the expectation of *g*:

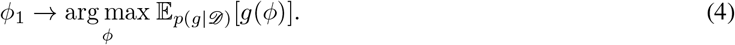

Next, we select a ‘challenger’, *ϕ*_2_, for which subjects’ preferences are most uncertain:

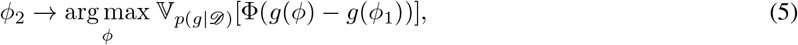

where 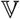 denotes the variance. Our algorithm is designed to balance exploitation (selecting values of *ϕ* that likely maximize *g*) and exploration (selecting values of *ϕ* for which the response is uncertain). In effect, the choice of champion encourages exploitation while the choice of challenger encourages exploration. In Fauvel and Chalk (2021a), we show that our Maximally Uncertain Challenge algorithm outperforms previous state-of-the-art PBO algorithms in a range of benchmark tests.

#### Limiting the search space

It was important to choose appropriate ranges for the optimization parameters, *ϕ*: too small, and the model would not be sufficiently flexible; too large and the optimization would be too difficult. To determine the range of values for the current diffusion and the axon sensitivity to current, we used the parameters inferred by Beyeler et al. (2019) from four patients. For the axon shape descriptors, we relied on the range of values determined by Jansonius et al. (2009) on 65 subjects. For the range of the remaining parameters (implant position and orientation), no data were available about postoperative implant movement at the time of the experiment, so we chose a realistic range of values. The bounds of the search space are summarized in table 1.

#### Kernel and hyperparameters selection

The performance of PBO algorithms is often strongly affected by the kernel and hyperparameters, which encode our prior assumptions about the latent function, *g*. We found that adapting the hyperparameters online, based on subjects’ responses, slowed down the algorithm and often led to overfitting. Because of this, we adopted a transfer learning strategy, where one participant who did not participate in the optimization experiment was asked to run 400 pairwise comparisons between random encoders, with two different perceptual models chosen at random. We then trained GP models on the training set, either with the Squared Exponential with Automatic Relevance Determination (SE-ARD), Matérn 3/2, or Matérn 5/2 kernel (see supplementary 4). We inferred the hyperparameters using type-II maximum-likelihood. We then tested the GP models on the test set and selected the best performing kernel by measuring each model’s Brier score (a widely used measure of probabilistic prediction accuracy). The best performing kernel (the Matérn 5/2 kernel) was later used in the experiment, using the hyperparameters learned on the training set. This can be seen as a way of transferring knowledge from previous patients to new ones (Supp Fig S3).

### 2.4 Psychophysics experiments

We presented simulated percepts generated by the *pulse*2*percept* model on a monitor at eye level. We only varied the amplitude of electrical stimulation, and not the pulse shapes or frequency, which were kept fixed at their default values (Beyeler et al., 2017a). We asked participants to sit approximately 50 cm from the screen, at which distance the simulated visual percepts, presented onscreen, covered 21°× 29° degrees of their visual field. Instructions were displayed on the monitor, and the participant answered using the computer keyboard. All participants had normal or corrected to normal vision and were between 22 and 67 years old (8 males and 16 females). One participant had prior experience in psychophysics experiments. Different optimization procedures were assigned in random order for each participant, to avoid the effects of fatigue, learning or attention lapses. Each optimization run included 60 iterations, and lasted approximately 8 minutes. 12 out of 24 subjects participated in the experiment remotely, using the remote access software TeamViewer (TeamViewer GmbH). We did not observe differences in the results between remote participants and participants who performed the tasks in the lab (Supp Fig S2).

#### Preference

To measure subjects’ preferences for different encoders after the optimization procedures, we simultaneously displayed the percepts predicted with two different encoders and asked the subject to indicate which one they preferred (**Fig 1B**). To get an accurate measure of subjects’ preferences we repeated the comparison for a set of 13 letters. The name of the letter was indicated above so that the subject knew what the reference stimulus was. The letters were selected either from the same set of 13 letters used during optimization (the ‘training set’) or from another set (the ‘test set’).

#### Visual acuity

We used two distinct tasks to measure visual acuity: the tumbling E, and the Snellen chart task. In the tumbling E task, we presented the letter E at various sizes at four different orientations and asked the subject to report the perceived orientation. In the Snellen chart task, we presented letters of various sizes and asked the subject to identify the letters (including the letters C, D, E, F, H, K, N, P, R, U, V, and Z).

We estimated participants’ visual acuity by fitting a psychometric function to their responses of the form:

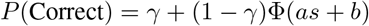

Where *s* is defined as the size of the finest detail in the stimulus (1/5 the size of the letter) in log of the visual angle in minutes of arc, *γ* is the chance level, and *a* and *b* are constants to be fitted. The visual acuity is defined as the inflection point of the curve,which corresponds to the letter size, *s*, for which 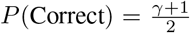. To efficiently measure *a* and *b*, we used the adaptive psychophysics procedure QUEST+ (Watson, 2017; Jones, 2018). The size of the visual field in the prosthetic vision simulations was 21°×29°, so the maximum size of the letter was 21°.

### 2.5 Bayesian data analysis

In order to compare the ability of different algorithms to optimize participants’ preference, we used the Bayes factor. For two algorithms *A*_1_ and *A*_2_ used to optimize encoders, we have the two hypotheses *H*_1_ (resp. *H*_2_): encoders found using *A*_1_ (resp. *A*_2_) are preferred to encoders found using *A*_2_ (resp. *A*_1_). The Bayes factor *B*_21_ is the ratio between the likelihoods of the two models: 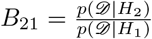. It gives an estimate of how strongly the data support *H*_2_ over *H*_1_.

To specifically adapt the analysis to the data-generating process in our experiments, we designed a method based on the Bayes factor, adapted from the extra-binomial variation detection method by Kass and Raftery (1995) (see also chapter 5 of Robert (2006)). An advantage of this analysis method (compared, e.g. to non-parametric tests) is that it models the noise in the preference measurement data.

For a participant *i*, the responses in a series of preference comparisons between two encoders follows a binomial distribution 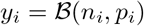, where *n*_*i*_ is the number of comparisons and *y*_*i*_ is the number of comparisons for which the participant reported that the encoder found using *A*_2_ is better than the one found using *A*_1_. We assumed the conjugate beta prior 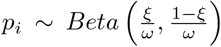. We assumed that *ω* is constant, and *ξ* follows a uniform distribution. Note that 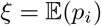. Under *H*_1_, the model is 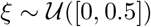 (meaning that on average, encoders derived using *A*_1_ are preferred to those derived using *A*_2_), whereas under *H*_2_, 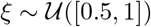.

The details of the computation of 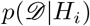 are described in supplementary 2. A *K* = log_10_ (Bayes factor) above 0.5, 1 or 2 corresponds to substantial, strong and decisive evidence in favor of *H*_2_ respectively. Conversely, a *K* = log_10_ (Bayes factor) below −0.5, −1 or −2 corresponds to substantial, strong and decisive evidence in favor of *H*_1_ respectively (Kass and Raftery, 1995).

Similarly, to compare visual acuity in the E and Snellen tasks between different encoders, we assumed that the differdifference in visual acuity ΔVA measured between encoders obtained with two methods *A* and *B* follows 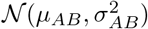. The two competing hypotheses are *H*_1_: *μ*_*AB*_ > 0 (that is, on average, *B* leads to encoders that correspond to better VA compared to *B*) and *H*_2_: *μ*_*AB*_ < 0. We infer 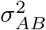 by measuring the empirical variance, and we put a uniform prior 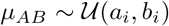, where *a*_1_ = *b*_2_ = 0, *b*_1_ = *v*, and *a*_2_ = −*v*, where *v* corresponds approximately to the maximum of visual acuity that we can measure. Again, the details of the computation of the likelihood 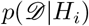 are described in supplementary 3.

## 3 Results

### 3.1 Model-based encoding of visual inputs

The standard approach in visual prosthetics is to down-sample the input image, and then linearly convert resulting brightness to current amplitudes or frequencies (Bloch and da Cruz, 2019). To implement this, we tessellated the input space so that each image pixel drove only the electrode closest to the pixel (Voronoi tessellation). The linear scaling was then adjusted so that the predicted percept would not saturate. In our simulations, using such an encoder lead to a highly blurred visual percept, where it was hard to distinguish the presented stimulus (**Fig 1C**, above).

If we knew the ‘true’ parameters, *ϕ*\*, describing how electrical stimulation is converted to a percept, then in theory we could directly infer the optimal encoder, ***W***_*ϕ*_*, that minimizes the visual distortion (Eqn 2). **Figure 1C** (below) shows the predicted visual percept after performing such an optimization. To compensate for current spread and axon activation, individual electrodes have a ‘center-surround receptive field’, where they are driven by bright pixels in the center and inhibited by bright pixels in the surround. The resulting visual percept is greatly improved, relative to the naive encoder that simply down-samples the image.

Unfortunately, however, we do not generally know the true perceptual model parameters, *ϕ*\*, for each subject. Nonetheless, our procedure implicitly restricts the high-dimensional set of possible encoders to a low-dimensional set of *optimal* encoders, ***W***_*ϕ*_. We were thus interested to see if we would still achieve performance improvements by using this low dimensional parameterization, even if we just chose *ϕ* randomly for each subject. To test this, we asked a subset of 14 participants to compare the perceptual quality of the naive encoder to a model-based encoder, ***W***_*ϕ*_, with random parameters, *ϕ* (chosen from the ranges shown in table 1; **Fig 2A-B**). We found that participants showed strong preference for the parameterized encoder, even with the parameters chosen at random (**Fig 2C**; log_10_ of the Bayes factor, *K* = 2.4, decisive evidence). Likewise, their visual acuity was significantly improved when using the model-based encoder compared to the naive encoder, in both the tumbling E task (log_10_ of the Bayes factor: *K* = −2.2, decisive evidence) and the Snellen task (*K* = −0.85, substantial evidence) (**Fig 2D**).

**Figure 2:**
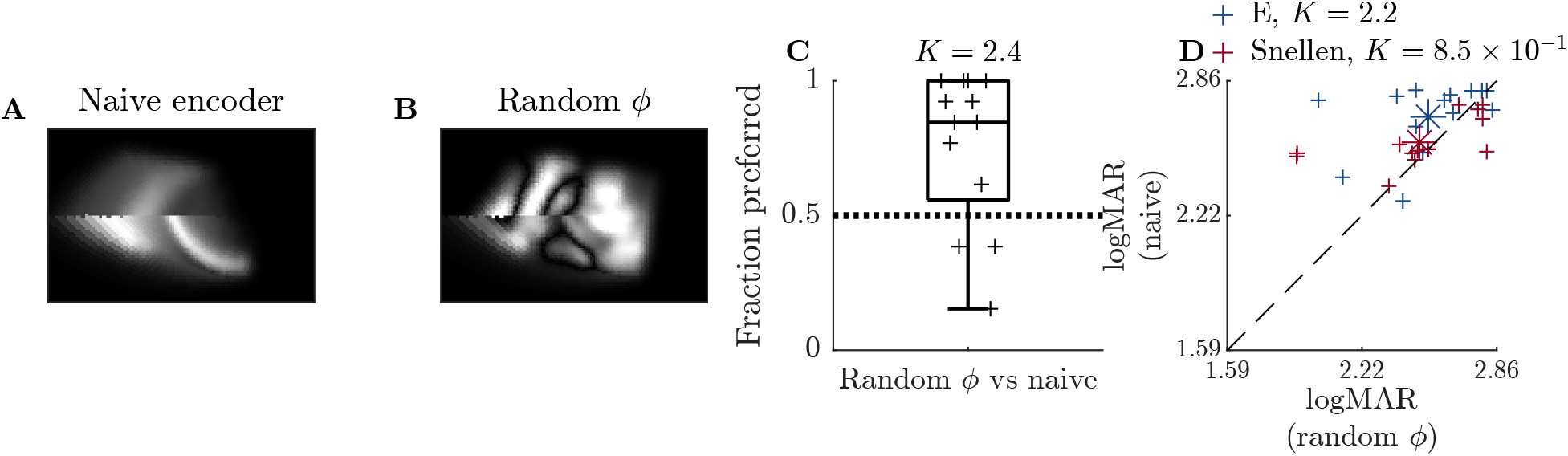
Effect of model-based parameterization of the encoder. (**A**) Percept predicted by the pulse2percept model using a naive encoder, where the pattern of electrode activation corresponds a down-sampled version of the image. The input stimulus is the letter ‘A’. (**B**) Predicted percept, using an encoder optimized with a random parameter, ϕ (we call this the ‘random-ϕ’ encoder). (**C**) Subjects’ preference for the naive encoder versus the ‘random-ϕ’ encode. In most cases participants prefer the random-ϕ (K = 2.4, N = 14, decisive evidence). (**D**) Visaul accuity (VA) measured in the tumbling-E (blue) and Snellen task (red) with the naive versus the random-ϕ encoder. The control encoder corresponds to better visual acuity in both the tumbling E (K = −2.2, N = 14, decisive evidence) and the Snellen task (K = −0.85, N = 14, substantial evidence). The mean of each distribution is denoted with asterisks.

### 3.2 Preference-based optimization of encoder

Next, we investigated the effect of optimizing the encoder, ***W***_*ϕ*_, based on subjects’ preferences. **Figure 3A** shows the duels presented to subjects on a typical experiment (left), along with percept generated using the inferred optimal encoder (right). The subject’s preferences on each trial are indicated by a red square. As data accumulates, the duels correspond to less distorted percepts, and the inferred percept also improves.

**Figure 3:**
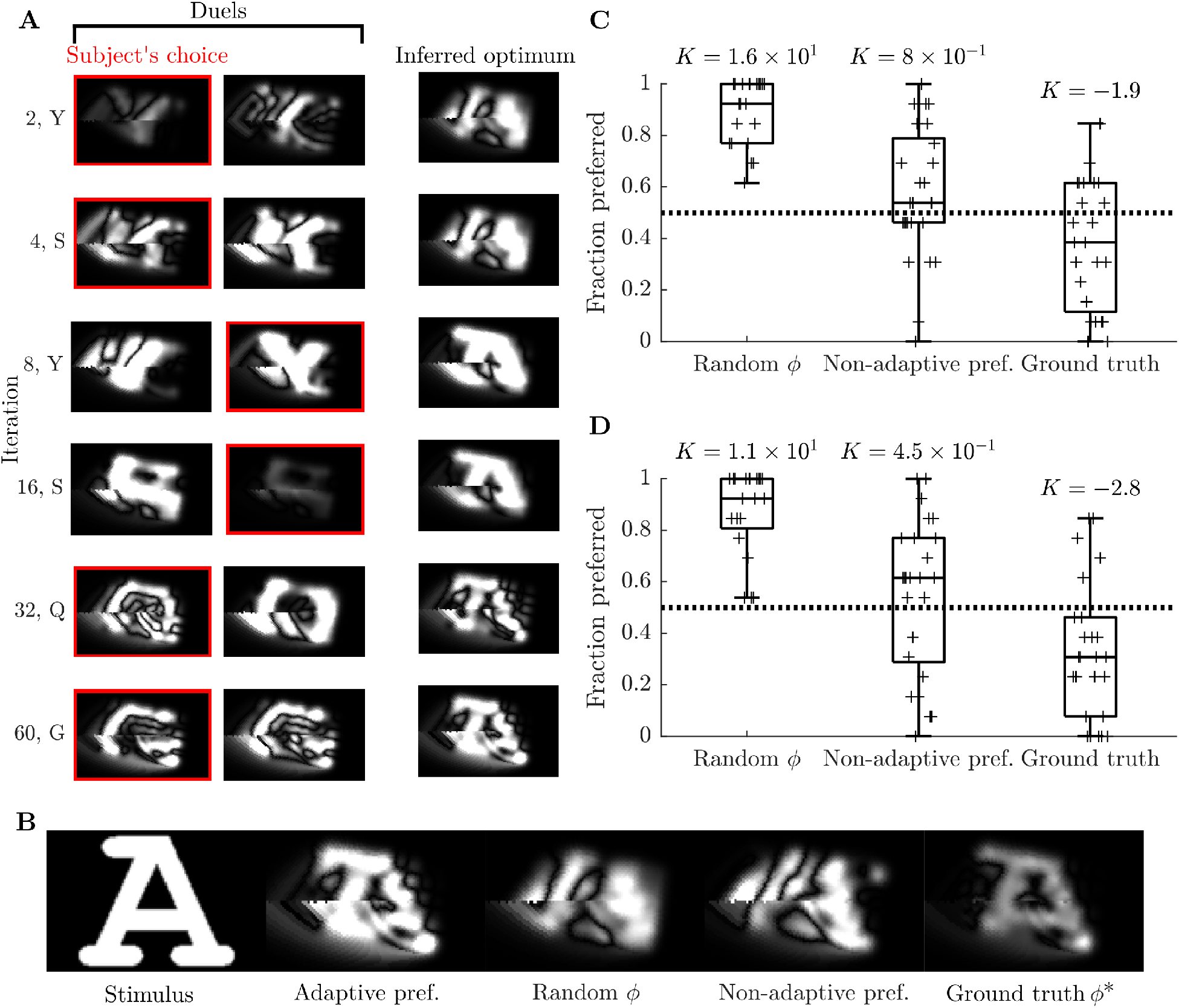
Results of preferential Bayesian optimization (PBO). (**A**) Illustration of a single optimization sequence. (left columns) Stimulus pairs (duels) presented to subjects on individual trials. The trial number and target letter identity are indicated on the left. Subjects’ preferences on each trial are indicated with a red outline. (far right) The percept predicted using the inferred optimal encoder at each iteration (with the letter A). As the optimization continues, the inferred optimal becomes less distorted. (**B**) Predicted percepts, after a single optimization run with: (i) adaptive PBO (‘adaptive pref.’; as in panel A); (ii) random duels during training (non-adaptive pref.); (ii) an encoder with random parameters (‘random ϕ’); and (iv) an encoder optimized using the ground truth parameters, ϕ* (‘ground truth’). The percepts tend to be discontinuous at the limit between the upper and lower hemifields, due to the geometry of the axons. (**C**) Subjects’ preferences for the encoder optimized using adaptive PBO, versus each of the controls. The encoder found using the adaptive procedure is preferred to the encoder with random parameters (K = 16, decisive evidence), and to the encoder obtained with the non-adaptive algorithm (K = 0.8, substantial evidence). However, the encoder based on the true model parameters (ground truth) is significantly preferred (K = −1.9, strong evidence). (**D**) Subjects preferences, evaluated using a new set of test stimuli (i.e. not used during the optimization process). While quality similar results were obtained, on this stimulus-set there is barely a noticeable difference between encoders derived from the adaptive or non-adaptive procedure (K = 0.45).

To assess the performance of our PBO algorithm, we compared it to: (i) a ‘true *ϕ*’ encoder, optimized using the ground truth parameters, *ϕ*\*; (ii) a ‘random *ϕ*’ encoder, optimized with random parameters, *ϕ*_random_ (as in the previous section); (iii) a non-adaptive algorithm, where the duels presented on each trial were generated using random parameters, (*ϕ*_1_, *ϕ*_2_). **Fig 3B** shows the resulting visual percepts after optimizing the encoder in one experiment. Here, the encoder inferred using the PBO algorithm (second left) results in a less distorted percept compared to the ‘random *ϕ*’ encoder (far right), or non-adaptive algorithm (second to left), though more distorted than with the ‘true *ϕ*’ encoder. These qualitative results were confirmed when we ran our optimization algorithm and controls on 24 subjects, each with a different simulated perceptual model, *ϕ*\*. After optimization, subjects were asked to say whether they preferred the PBO encoder or the controls on 13 randomly chosen letters (**Fig 3C**). Overall, the PBO encoder was significantly preferred relative to the ‘random *ϕ*’ (*K* = 16, decisive evidence) and encoders found using the non-adaptive optimization (*K* = 0.80, substantial evidence), but worse than the ‘true *ϕ*’ encoder (*K* = −1.9, strong evidence). Qualitatively similar results were obtained when we assessed subjects’ preferences using a different set of letters, that had not been presented during training (**Fig 3D**).

### 3.3 Transfer to other tasks

To be useful to patients, it is important that our optimization procedure generalizes to improve subjects’ performance in different tasks. To assess this, we assessed subjects’ performance in two tasks commonly used to measure visual acuity: the ‘tumbling E’ task (**Fig 4A**), where the goal is to detect the orientation of a square E among four possibilities, and the ‘Snellen chart’ task (**Fig 4B**), where the subject has to recognize a letter. We measured a significant improvement in visual acuity with the PBO algorithm, compared to encoders with ‘random *ϕ*’ encoders (**Fig 4C**), in both the tumbling-E (*K* = 4.9, decisive evidence) and Snellen tasks (*K* = 3.0, decisive evidence). Performance was slightly better in the tumbling E task, compared to the non-adaptive algorithm (**Fig 4D**, *K* = 0.54, substantial evidence), although we saw no significant difference for the Snellen task. Consistent with our previous results, the optimized encoder performed worse than the ‘true *ϕ*’ encoder (*K* = −4.8 and *K* = −2.2, decisive evidence) in both tasks.

**Figure 4:**
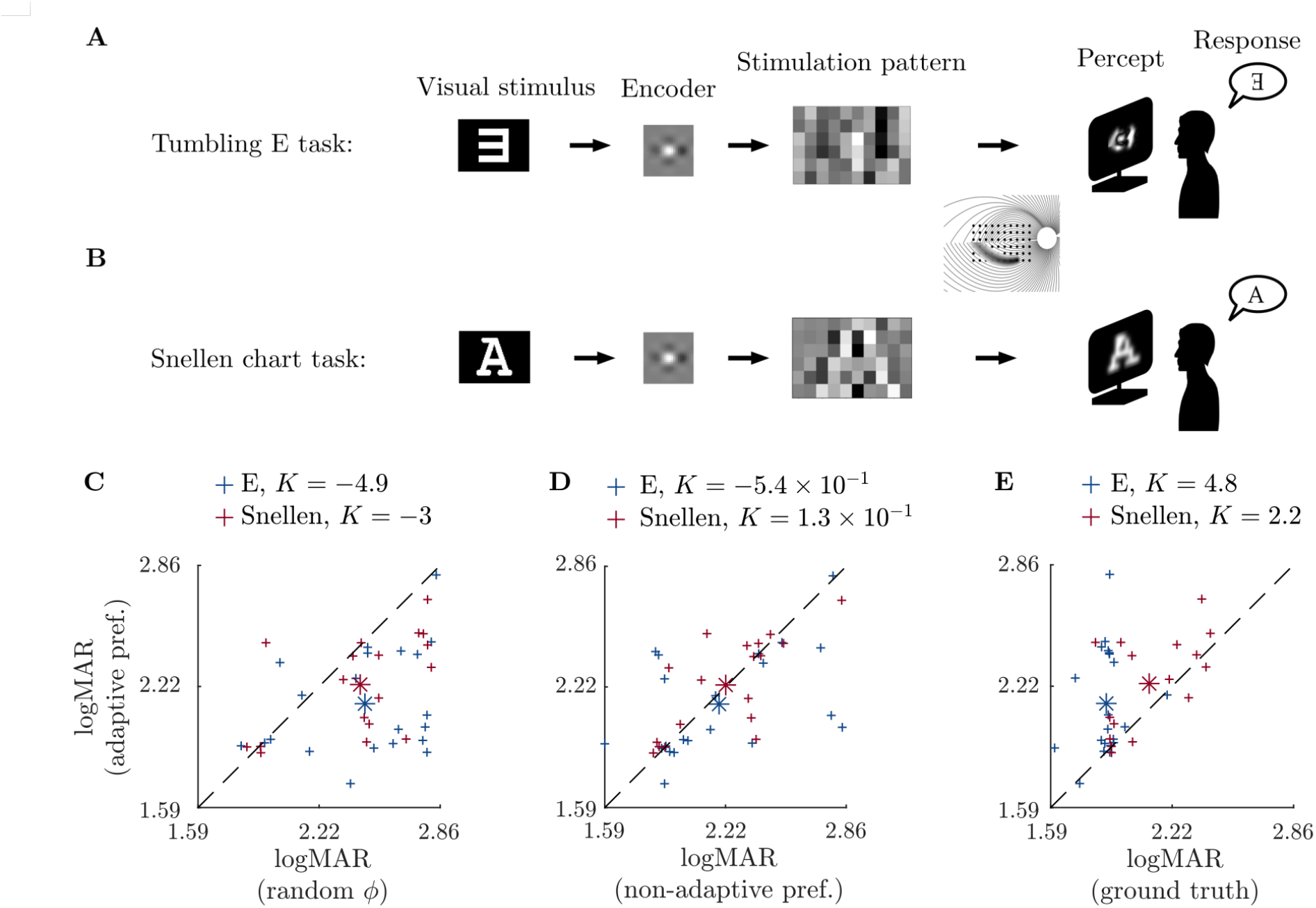
Visual acuity measurements following preference-based optimization (PBO). (**A**) The tumbling E task consists in identifying the orientation of a letter E among four possibilities. (**B**) The Snellen chart task consists in recognizing a letter from the Snellen letters chart. We measured visual acuity (VA) with a specific encoder in each of these tasks by varying the letter size. (**C**) Encoders found using adaptive PBO (adaptive pref.) lead to better visual acuity compared to encoders with random parameters (random ϕ) in both the tumbling E (K = 4.9, decisive evidence) and Snellen chart task (K = 3, decisive evidence). Asterisks represent the centroids. (**D**) There is only a slight improvement in the tumbling E task with adaptive PBO, compared to encoders non-adaptive PBO (non-adaptive pref., K = 0.54, substantial evidence), and no significant difference in the Snellen chart task (K = −0.13). (**E**) The encoders found using adaptive PBO do not match the performance of the encoders based on the true model (ground truth) in both the tumbling E task (K = −4.8, decisive evidence) and Snellen chart task (K = −2.2, decisive evidence).

### 3.4 Performance-based optimization

An alternative strategy would be to optimize the encoder so as to maximize subjects’ performance in a specific visual task. To test the efficacy of such a strategy, we ran a second set of experiments in which we tried to optimize subjects’ performance in a tumbling E task. For these experiments, a letter size covering 7.4°of visual field was chosen in pilot studies, so as to achieve a success rate of about 50% using random *ϕ* encoders. The optimization framework was the same, as both preference and performance in forced-choice tasks can be modeled using binary classification models. As is common in binary Bayesian optimization, we used a Thompson Sampling algorithm (Thompson, 1933) to select the encoder on each trial (see Fauvel and Chalk (2021a) for implementation details).

Performance-based optimization did not lead to improved visual acuity compared to ‘random *ϕ*’ encoders (**Fig 5A**). Preference-based optimization procedure led to a improved visual acuity (**Fig 5B**) in both tumbling E (*K* = −3, decisive evidence) and Snellen chart tasks (*K* = −3.1, decisive evidence) compared to performance-based optimization. Finally encoders optimized using preference-based learning were strongly preferred to encoders optimized using performance-based learning (**Fig 5C**; *K* = −7.8, decisive evidence). Indeed, performance-based encoders were not preferred to ‘random *ϕ*’ encoders (*K* = 0.1).

**Figure 5:**
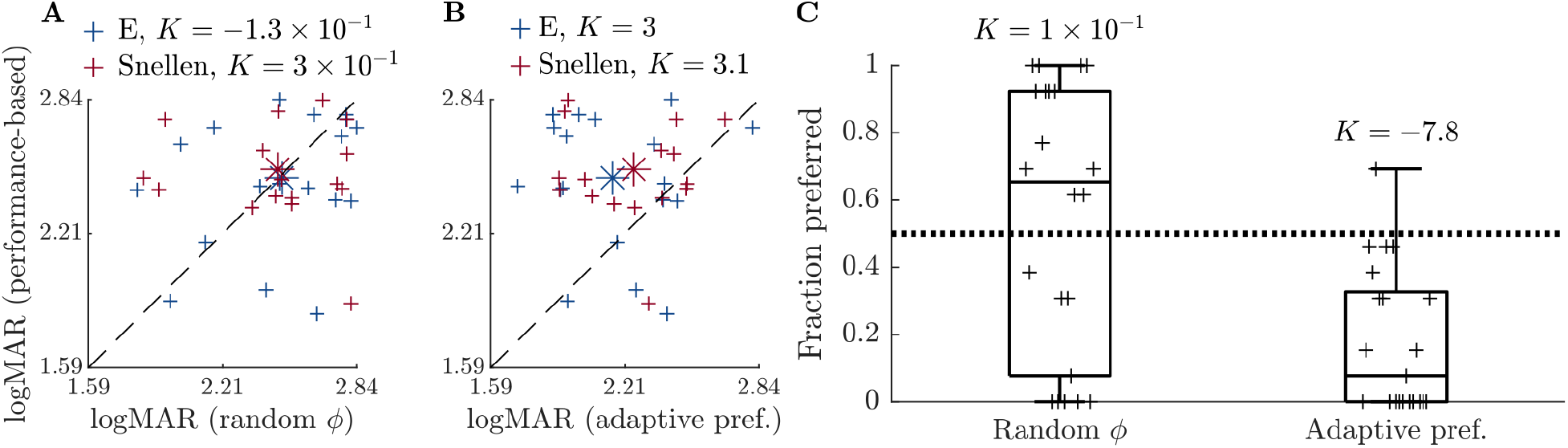
Performance-based optimization. (**A**) Comparison between visual acuity measured with the encoders found with performance-based optimization and encoders with random parameters (random ϕ). There is no significant improvement after optimization in the tumbling E task (K = 1.3 × 10^−1^) or Snellen task (K = −3.0 × 10^−1^). Asterisks represent the centroids. (**B**) Preference-based encoders obtained with adaptive PBO (adaptive pref.) lead to better visual acuity in both the tumbling E (K = −3.0, decisive evidence) and Snellen task (K = −3.1, decisive evidence). (**C**) Performance-based encoders are not significantly preferred to encoders with random parameters (K = 0.10). Preference-based encoders are preferred to performance based-encoders on the set of stimuli that was not used to optimize the encoders (K = −7.38, decisive evidence).

### 3.5 Robustness to model mismatch

Thus far we assumed that subject’s percepts are exactly described by the *pulse2percept* model, albeit with unknown parameters, *ϕ*\*. With real patients this may not be the case; there may be some mismatch between the assumed perceptual model, and what patients actually observe. To test whether our procedure is robust to such model misspecification, we ran our experiments using fixed, misspecified parameters to describe the paths of axons in the retina (**Fig 6A-B**). We first confirmed that the misspecification of the encoder led to severely degraded percepts (**Fig 6C**), resulting in poor performance in both the tumbling E and Snellen tests of visual acuity (**Fig 6D**; *K* = −8.6 and *K* = −13 for the E and Snellen task respectively, decisive evidence).

**Figure 6:**
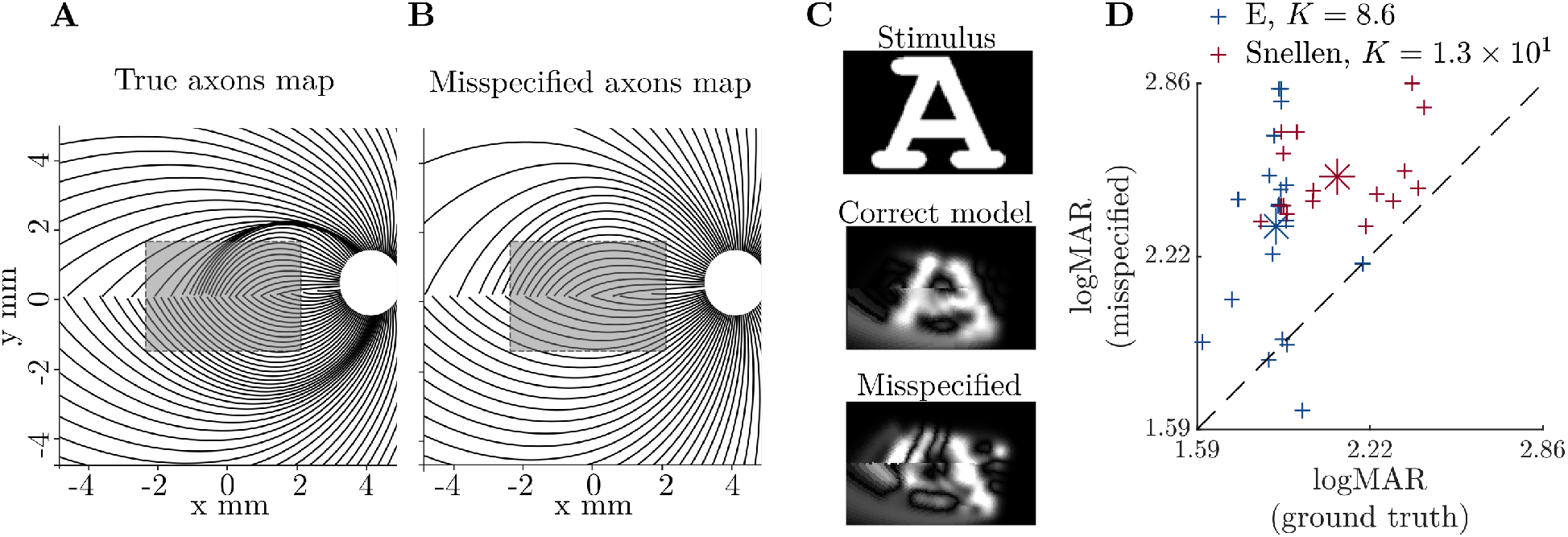
Mdel misspecification. (**A**) Axon map in the true patient model. The shaded area corresponds to the location of the implant. (**B**) Axon map in the misspecified model. Note how the axon trajectories are affected, in particular in the region below the implant. (**C**) Example of percept predicted with the encoder derived from the true perceptual model (correct model) or with the encoder derived from the misspecified perceptual model, where all parameters were set at their true value, except the two axon shape descriptors (misspecified). Note that the percept quality is severely affected by the model misspecification. (**D**) Effect of axon shape descriptors misspecification on visual acuity. With all parameters at their true value, except the two axon shape descriptors, visual acuity is significantly reduced in both the Snellen chart task (K = −13, decisive evidence) and the tumbling E task (K = −8.6, decisive evidence) compared to the encoder based on the ground truth parameters. The asterisks represent the centroids.

Optimizing the encoder with the misspecified model led to encoders that were preferred compared to the ones corresponding to random parameters and a misspecified model (*K* = 1.5, strong evidence), as well as to the ‘random *ϕ*’ encoders (*K* = 1.6, strong evidence) (**Fig 7B**). This suggests our optimization procedure is robust to model misspecification. However, the optimized encoder was not preferred compared to an encoder with all parameters but the axons shape descriptors at their true value (**Fig 7B**), suggesting our algorithm was not able to fully compensate for the model misspecification. On average, performance was also improved by the PBO algorithm, relative to the ‘random *ϕ*’ encoder, in both the tumbling E task (*K* = 1.8, strong evidence) and the Snellen chart task (*K* = 0.55, substantial evidence), despite model misspecification (**Fig 7C**).

**Figure 7:**
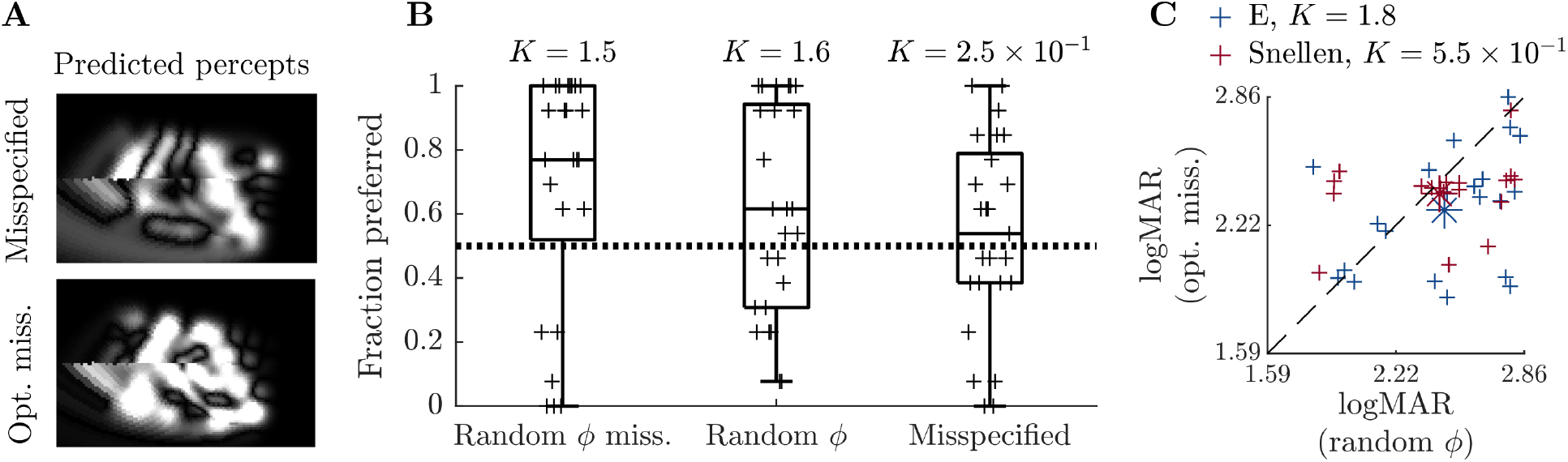
Robustness of optimization to model misspecification. (**A**) Example of percepts predicted for the stimulus in figure 6C, with (above) an encoder optimized using the misspecified model or (below) an encoder optimized using PBO with a misspecified model. Here, the optimization procedure found an encoder that partly compensated for the misspecification. (**B**) Encoders optimized using a misspecified model were preferred to encoders where all parameters were selected at random except the misspecified axon shape descriptors (random ϕ miss., K = 1.5, strong evidence), and encoders with all parameters random (random ϕ, K = 1.6, strong evidence). However, these encoders were not preferred on average to encoders found by inverting the misspecified model (misspecified, K = 0.25). This shows that despite the robustness of the optimization procedure, the limited encoder flexibility could not fully compensate for the misspecification. (**C**) Visual acuity was better with encoders optimized based on a misspecified model (opt. miss.) compared to random encoders in both the tumbling E task (K = 1.8, strong evidence) and the Snellen chart task (K = 0.55, substantial evidence). The asterisks represent the centroids.

## 4 Discussion

We provided a proof of principle for a new human-in-the-loop (HILO) optimization strategy to tune retina implant encoders. HILO is limited by the fact that we only have access to limited noisy data, which takes time to acquire. Moreover, the choice of optimization criteria that could lead to general functional vision improvements is challenging. To overcome these challenges, our proposed strategy is based on two ideas: first, we use a model of prosthetic vision to reduce the dimensionality of the encoder; second, we optimize the encoder using patients’ subjective reports through pairwise comparisons between encoders. We assessed our strategy on healthy subjects using a state-of-the-art prosthetic vision simulator (Beyeler et al., 2017a). Our method is readily applicable to real-life patients, and would solely require implementing our algorithm in the visual processing unit.

### 4.1 Model-based percept shaping

Current spread in the retina can result in cross-talk between electrodes, and overlapping phosphenes (**Fig 1A**). Most previous work considered this cross-talk to be detrimental and sought to reduce it (Lorach et al., 2015). Recently however, Spencer et al. (2018) showed, using a model describing how prosthetic stimulation activates RGCs, how to leverage cross-talk between electrodes to optimize the stimulation protocol, and control the pattern of RGC activity. Similar strategies were applied for transcranial direct current stimulation, so as to achieve intense and focal stimulation (Dmochowski et al., 2011). Chen et al. (2020) also followed a very similar approach for retinal prostheses, but instead of optimizing the stimulation based on an estimate of the tissue response, the optimization was based on the electrodes induced electric fields.

Our approach differs to these papers, as we focused on directly shaping the perceived phosphenes, using a model of prosthetic vision, and we further optimized the encoder based on patients’ feedback. We show how cross-talk between electrodes can be leveraged to compensate for the perceptual distortion due to activation of passing axons and current spread (**Fig 1A-B**).

### 4.2 Closed-loop optimization

A potential advantage of our closed-loop approach is that we could adjust the encoder to deal with the considerable differences in the perceptual experiences of different patients (Beyeler et al., 2019). In contrast, previous work, where the encoder was optimized in open-loop to improve patients’ performance in a given visual task (such as reading) (Feng and McCarthy, 2013; Luo and da Cruz, 2016; Barnes et al., 2016; Han et al., 2021), would not be able to take into account such differences between patients.

A perception-based closed-loop optimization of retinal implants was proposed in a series of papers by Eckmiller et al. (1999); Becker et al. (1999); Eckmiller et al. (2005, 2007). However, certain aspects of their approach would be hard to realize on real patients. For example, in their work subjects were asked to perform a comparison between a predicted percept and a simultaneously presented undistorted ‘target’ image (such as a white circle on a dark background). Real patients, however, will not have access to such an undistorted target image, and it is thus hard to see how this procedure could be replicated on them. Our approach, on the contrary, is based on a comparison between percepts produced by different encoders, and the reference to the cue is purely symbolic (we displayed the name of the letter corresponding to the stimulus, but in real-life patients the letter could be indicated orally).

Further, Becker et al. (1999) and Eckmiller et al. (2007) relied on genetic and reinforcement learning algorithms, which are known to be less data-efficient than Bayesian Optimization (Turner et al., 2021). Consistent with this, the training procedure in the work of Becker et al. (1999) and Eckmiller et al. (2007) took 60-120 minutes to converge, while in our work significant performance improvements were achieved in several minutes. More importantly, and in contrast to our work (**Fig 1D** & **Fig 2**), Becker et al. (1999) and Eckmiller et al. (2007) did not demonstrate that their procedure generalized to improve subjects’ perception of different shapes other than those used during training.

Another line of research seeks to optimize the stimulation protocol so as to mimic the activity of retinal neurons under natural conditions (Nirenberg and Pandarinath, 2012; Twyford et al., 2014; Guo et al., 2018; Lee and Im, 2019; Shah et al., 2019). This approach is based on the hypothesis that we have access to a metric relating RGCs activity to visual function (Shah et al., 2017), typically assuming that the more natural the neural response is, the better. In contrast, in our work, we do not explicitly assume any metric defining which patterns of neural activity are ‘best’, and instead rely on subjects to directly state what they prefer.

Indeed, our results support the idea that there may be no simple metric to define *a priori*, which patterns of stimulation are preferred by subjects. For example, an obvious choice of metric is the mean-squared-error (MSE) between the predicted percept and the original stimulus. Surprisingly, however, in our experiment we found that the MSE was not smaller for optimized encoders, compared to encoders with random parameters (on the contrary, it was significantly larger; Supp Fig S4A). Thus, it seems that the value subjects attribute to a specific encoder cannot be predicted by the corresponding MSE, and thus, their feedback (about which encoders they prefer) may be required to optimize the encoder.

Another disadvantage of previous approaches to closed-loop stimulation optimization in retinal implants is that demonstration of performance usually relies on an assumed known decoder (i.e., a mapping from stimulation patterns to percepts, see e.g. Nirenberg and Pandarinath (2012); Eckmiller et al. (2005)). As a consequence, the results merely correspond to an upper bound on improvement, and it is not clear whether there would be effective improvement in case of encoder-decoder mismatch. In our work, on the other hand, the model is only used to reduce the number of parameters. It is not an essential part of the optimization process. As a result, while misspecifying the model can slightly reduce performance, it does not overly hinder the optimisation process (**Fig 7**). This suggests that our method could be applied in real-life patients even if there is a severe mismatch between the model used to parameterize the encoder and the true perceptual model. However, our results show that optimizing the parameters does not fully compensate for the misspecification, so that the device does not reach its full vision restoration potential. This is due to the fact that by parameterizing the encoder, we restrict the search of the optimum to a subspace in the space of all possible encoders. This makes the optimization easier but limits flexibility.

### 4.3 Preference-based optimization

We used preferential Bayesian optimization (PBO) to optimize the encoder, an efficient method that builds a model of the subject’s preferences to adaptively select new configurations to test. This framework leverages prior information from other patients: in our experiments, the search space was constrained by the knowledge of the distribution of the perceptual model parameters among patients, while the hyperparameters of the GP kernel were learned on one participant and then used for the others. We showed that preferential Bayesian optimization of the encoder led to significant improvement in perceptual experience compared to randomly selected encoder parameters. Notably, the adaptive optimization procedure led to better results than the non-adaptive one.

PBO was first applied to procedural animation design (Brochu et al., 2010). Later, it was used to tune hearing aids (Nielsen, 2015). To solve the problem of generalizing improvement across many conditions, Kupcsik et al. (2018) applied PBO to optimize a robot-to-human object handover across a variety of randomly chosen contexts (objects size and type) based on human feedback. The idea that preference measures may be more relevant than objective performance is further supported by the work of Tucker et al. (2019), who applied PBO to the problem of optimizing two gait parameters of the Atalante lower-body exoskeleton. The goal was to maximize user comfort when walking inside the exoskeleton. In this paper, interestingly, no correlation was found between metabolic expenditure and user preferences.

To our knowledge, the first application of PBO to a brain-computer interface was reported by Lorenz et al. (2019), who used it to maximize the phosphenes brightness elicited in transcranial alternating current stimulation when tuning current frequency and phase. Recently, Zhao et al. (2021) applied PBO to epidural spinal cord stimulation. However, in none of the aforementioned studies did the authors compare their acquisition rule (the criterion used to select new samples) to, for example, random sampling.

### 4.4 Preference- versus performance-based optimization

Our rationale for using preference-based optimization came from the idea that it allows to take into account and balance many different aspects of perception, that are difficult to assess by measuring subjects’ performance in any given task. However, it was possible that preference-based optimization might lead to percepts that the subject prefers (for their own, subjective, reason), but which do not improve general visual function. In reality, our results argue against this, as preference-based optimization improved performance on different stimuli (Fig 3D), and different tasks (Fig 4).

In contrast, while performance-based optimization may appear more ‘objective’, a general problem is that improvement in a specific task, or set of tasks, may not be related to vision improvement in general. Indeed, the optimization process may emphasize artefactual perceptual cues that are not related to vision. This problem is related to the more general vision testing problem in the field of vision restoration and has been extensively discussed by Peli (2020). As a consequence, we expected preference-based optimization to generalize better than a single performance-based optimization. However, we could not test this hypothesis as the performance-based optimization we assessed did not improve subjects’ visual function.

Multiple reasons may explain that the performance-based optimization failed. First, performance measurements in binary tasks may be less informative than pairwise comparisons. Second, the task may be either too difficult or too easy so that responses are not informative. If the encoder improves over the course of the optimization process, without varying the difficulty of the task, the optimization algorithm will not differentiate between encoders that are equally good for that level of difficulty. At the time we did the experiments, we could not find existing optimization algorithms addressing the problem of tuning the task difficulty (but see Fauvel and Chalk (2021b)).

Preference-based optimization can suffer from a similar drawback. For a given set of stimuli, the subject may find all the encoders that are proposed to him equally good or equally bad. Again, this highlights the importance of choosing appropriate stimuli. In our experiment, however, we did not have to hand-tune the set of stimuli. This suggests that designing a preference-based optimization protocol is much easier than selecting forced-choice visual tasks. Finally, participants frequently reported they found the preference judgment task much more engaging, which is of practical importance for patients who have to engage in prolonged rehabilitation protocols.

### 4.5 Limits and future developments

Likely, more accurate prosthetic vision models will also include more parameters. While this could improve the extent to which visual function can be improved thanks to increased encoder flexibility, this would also make the optimization problem more difficult. In general, for a given amount of data, the encoder dimension determines a tradeoff between the encoder’s ability to flexibly adapt to the patient’s preference, and the difficulty to learn this preference as dimension increases. In particular, highly flexible encoders would require enhancing the Bayesian optimization algorithm with features specifically designed for high-dimensional optimization (see e.g. Gardner et al. (2017); Wang et al. (2017); Mutný and Krause (2018); Rolland et al. (2018); Li et al. (2017); Zhang et al. (2019)).

To limit the total duration of the experiment for individual participants, we restricted optimization sequences to 60 iterations. At each iteration, computing the new candidate encoders required to run the *pulse*2*percept* model, which took a few seconds, so that an optimization sequence lasted about 8 minutes. Our results show that despite considerable improvement in encoder quality, this was not sufficient to converge to the preferred encoder. In application to real-life patients, an optimization sequence could probably last a few hours, allowing the encoder to improve even further.

We did not consider the temporal dynamics of stimulation and percepts. Electrical stimulation induces a desensitizing phenomenon in RGCs, whereby phosphenes gradually fade over time (Horsager et al., 2009; Freeman et al., 2011; Pérez Fornos et al., 2012). Our approach could generalize to include the temporal dimension in the encoder and in the tasks. For example, we could ask the subjects to compare their perceptual experience between encoders for static stimuli exactly as we did, but the subjects would also have to consider the temporal dynamics of their percept (for example, a subject would likely prefer a percept that does not fade to one that does). Our approach could also be used to tune stimulation parameters other than currents amplitudes, such as pulses shapes and frequency, for more precise tuning.

Our model-based parameterization of the retinal prosthetic encoder corresponds to minimizing the expected distance between the stimulus and the percept (see equation 2). In practice, however, the distortion minimization problem is constrained due to the charge injection limit of metallic electrodes. The Argus II Surgeon Manual (Second Sight Medical Products: Argus II Retinal Prosthesis System Surgeon Manual (2013)) specifies that the maximum current per electrode is 1.0 mA. However, the *pulse2percept* model predicts that the percept brightness scales approximately linearly with the pulse frequency (Nanduri et al., 2012). That way, it is possible to reduce pulse amplitudes while increasing the frequency to stay below safety limits. Another, more general approach to enforce such constraints on the encoder would be to use a quadratic optimization algorithm with linear constraints.

In general, for more complex encoders including varying pulse shapes, frequencies, etc, which involve nonlinear effects, more advanced optimization techniques would be required to invert the perceptual model. Note, however, that this is fully compatible with our approach, the only difference being the time taken to compute a given encoder.

In our experiments, the preference-learning task was general enough so that the optimized encoder was appropriate in a range of different tasks. A possible implementation would specifically adapt the encoder to different activities or environments. One could also consider adaptively varying the stimuli that are presented during the optimization sequence, to incorporate more complex stimuli with finer detail. Hopefully, this would allow the user to make refined comparisons, and enhance the transfer of improvement to other tasks.

A significant advantage of our approach is that subjects could optimize their retina implant encoder on their own during daily life, without expert assistance. This could be particularly useful as the retina continues to degenerate over time, the implant/retina interface evolves, and the subject learns to make use of prosthetic vision (Beyeler et al., 2017b). The long-term influence of artificial stimulation of the retina on network architecture is still unknown and is a remaining concern, but our system could circumvent those changes by adapting the way the electrode array transmits visual stimuli.

## Code availability

The Matlab code to perform the experiments and run the analyses is available at https://github.com/TristanFauvel/Retinal_Prosthetic_Optimization_Code.git. The implementation of the QUEST+ adaptive psychophysics procedure is from Jones (2018). To display stimuli and record responses, we used the Matlab toolbox Psychtoolbox (Brainard, 1997; Pelli, 1997; Kleiner et al., 2007)

## Acknowledgments

We thank Olivier Marre for fruitful discussions.

This research was supported by the Institut National de la Santé et de la Recherche Médicale and the Centre Hospitalier National des Quinze-Vingts.

## Competing interests

The authors have no competing interests to declare.

## Ethics

All experimental protocols were approved by the Comité de protection des personnes of the Institut National de la Santé et de la Recherche Médicale.

## Contributions

Tristan Fauvel: Conceptualization, Methodology, Software, Validation, Formal Analysis, Investigation, Data Curation, Writing - original draft preparation, Writing - review & editing, Visualization.

Matthew Chalk: Conceptualization, Methodology, Software, Resources, Writing - original draft preparation, Writing - review & editing, Supervision, Project Administration, Funding Acquisition.

## Supplementary material

### 1 Supplementary figures

**Figure S1:**
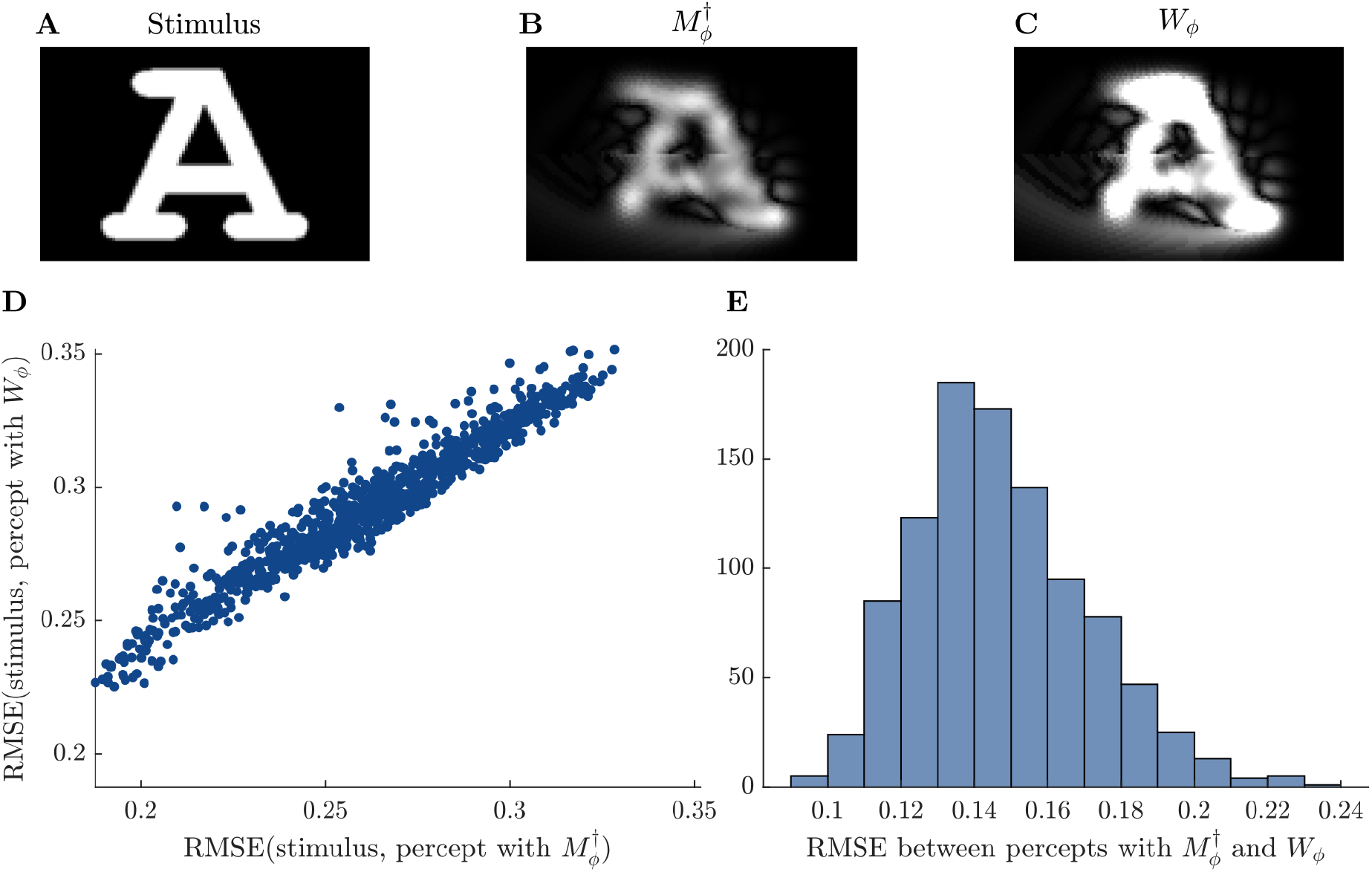
Comparisons between percepts predicted by the pulse2percept model for the stimulus in **A**, either obtained with the analytical solution of the simplified problem 2 (in **B**), 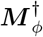, or the encoder that is a numerical solution of the exact minimization problem **W**_ϕ_ (in **B**, note that we specifically optimized the amplitudes of the 60 electrodes for this stimulus). These two solutions lead to virtually identical percepts. **D**. Comparison of the distance (RMSE) between the stimulus and the percept obtained using either 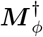 or **W**_ϕ_ for 1000 randomly chosen perceptual models. **C**.Histogram of the RMSE between percepts obtained either with 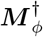 or **W**_ϕ_ for 1000 randomly chosen perceptual models.

**Figure S2:**
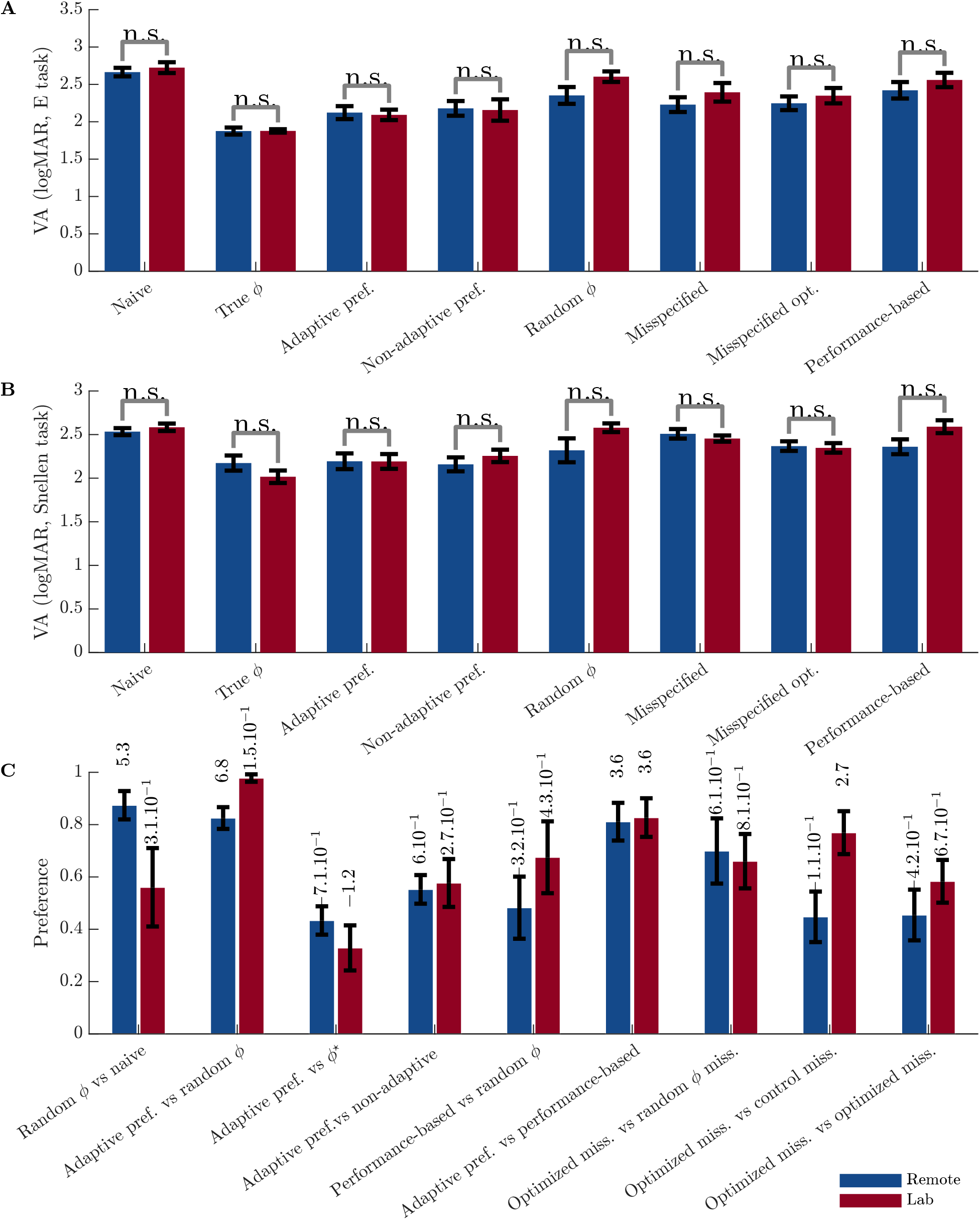
Comparison of results for participants who participated to the experiment remotely (in blue) or in the lab (in red). **A**. Visual acuities measured in the tumbling E task. **B**. Visual acuities measured in the Snellen task. Visual acuities do not differ significantly between groups (two-samples t-test at 0.05 significance threshold) **C**. Preference comparisons between encoders. There is no contradiction between the two groups.

### 2 Bayesian analysis of preference data

In order to compare preference between encoders, we used the following Bayesian analysis method adapted from Kass and Raftery (1995) (see also chapter 5 of Robert (2006)): For two algorithms *A*_1_ and *A*_2_ used to derive encoders, we have the two hypotheses *H*_1_ (resp. *H*_2_): encoders found using *A*_1_ (resp. *A*_2_) are preferred to encoders found using *A*_2_ (resp. *A*_1_). For a participant *i*, the responses in a series of preference comparisons between two encoders follows a binomial distribution 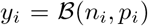, where *n*_*i*_ is the number of comparisons and *y*_*i*_ is the number of comparisons for which the participant reported that *A*_2_ is better than *A*_1_. We put the conjugate beta prior 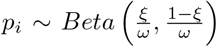. Note that 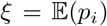. We assume that *ω* is constant, and *ξ* follows a uniform distribution. Under *H*_1_, the model is 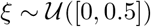, whereas under 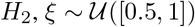.

The Bayes factor *B*_21_ is the ratio between the likelihoods of the two models: 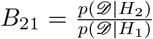. It gives an estimate of how strongly the data support *H*_2_ over *H*_1_.

We have:

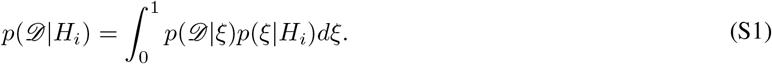

So for *H*_1_, by noting *G* the number of participants:

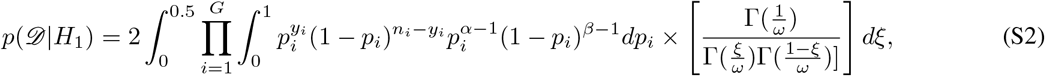

where 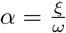 and 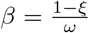.

So:

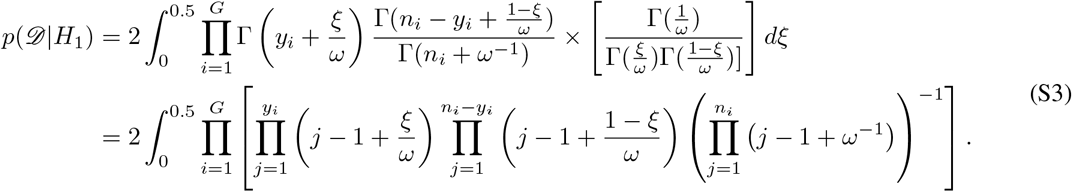

So the integrand in the likelihood is simply a polynomial function of *ξ* that can easily be integrated. Similarly, for the alternative hypothesis *H*_2_:

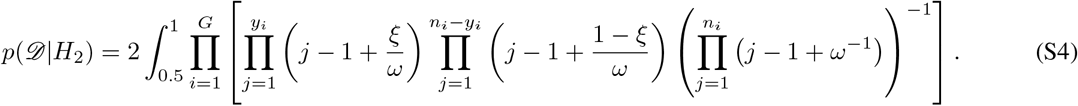

For each comparison between algorithms, *ω* was estimated empirically, by noting that: 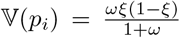. So we computed the empirical mean of the 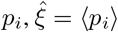 and the empirical variance 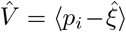, and solved for 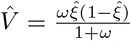. We then evaluated the integral numerically. An advantage of this analysis method is that it models the noise in the preference measurement data.

### 3 Bayesian analysis of visual acuity data

We have:

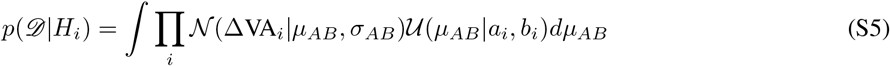

Based on S5, the Bayes factor can be computed via Monte-Carlo integration.

### 4 Kernel definitions

In the following definitions, *ρ* is the lengthscale of the kernels (positive constant), and *σ* is a positive scaling parameter.

The Matérn kernels is defined as:

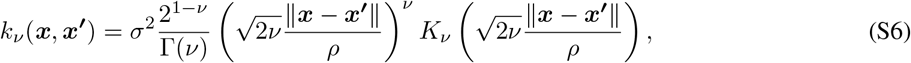

where Γ is the gamma function, *K*_*ν*_ is the modified Bessel function of the second kind and *ν* is a positive parameter. For example, the Matérn 5/2 kernel is defined as:

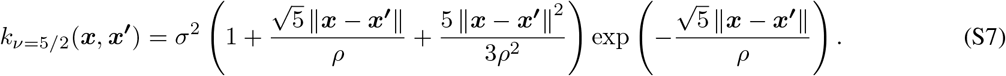

The Squared-Exponential kernel is defined as:

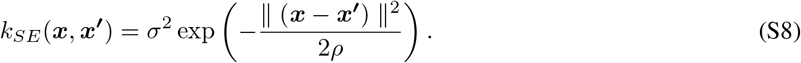

For stationary local kernels such as the Matérn and the Squared-Exponential kernels, it is possible to specify a different lengthscale for each input dimension. This is known as Automatic Relevance Determination (ARD). For example, the SE-ARD kernel is defined as:

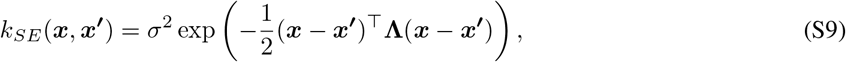

where **Λ** = diag(*λ*_1_, … , *λ*_*D*_). *D* is the dimension of the input space and the *λ_i_* are positive constants.

### 5 Instructions to the participants

The exact instructions given to the participants were the following (eventually translated to French for native French speakers):

- Preference comparisons: ‘The image corresponds to a distorted version of a letter (printed above), indicate which one you find the most recognizable or least distorted using the left or right arrows.’
- Tumbling E task: ‘The image corresponds to a distorted version of a letter E, which can take 4 different orientations. Indicate the orientation by pressing the corresponding arrow keys.’
- Snellen chart task: ‘The image corresponds to a distorted version of a letter. Press the keyboard to identify the letter.’

### 6 Transfer learning

**Figure S3:**
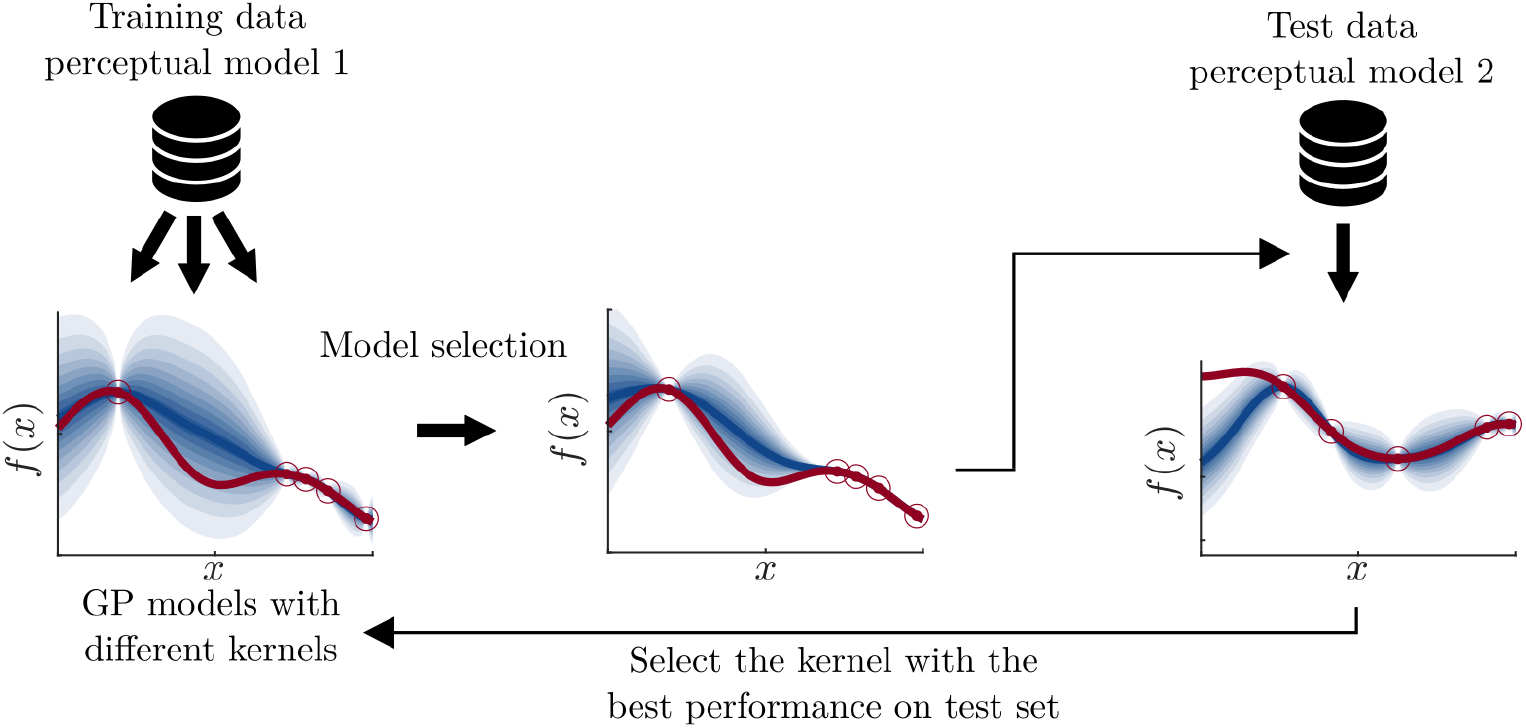
Illustration of the principle of hyperparameters transfer. The blue areas correspond to the GP posterior distribution. Red circles correspond to data points, and the red line corresponds to the true function. Two datasets of 400 comparisons between encoders are built using two different perceptual models. GP models are trained on the training dataset with three different kernels, and for each kernel, the hyperparameters are inferred using type-II maximum likelihood. After this model selection step, GP models are learned using the same hyperparameters on the test data set. The kernel for which the Brier score is the highest on this second data set is used in the experiment, with the same hyperparameters.

### 7 Relationship between MSE and preference between encoders

**Figure S4:**
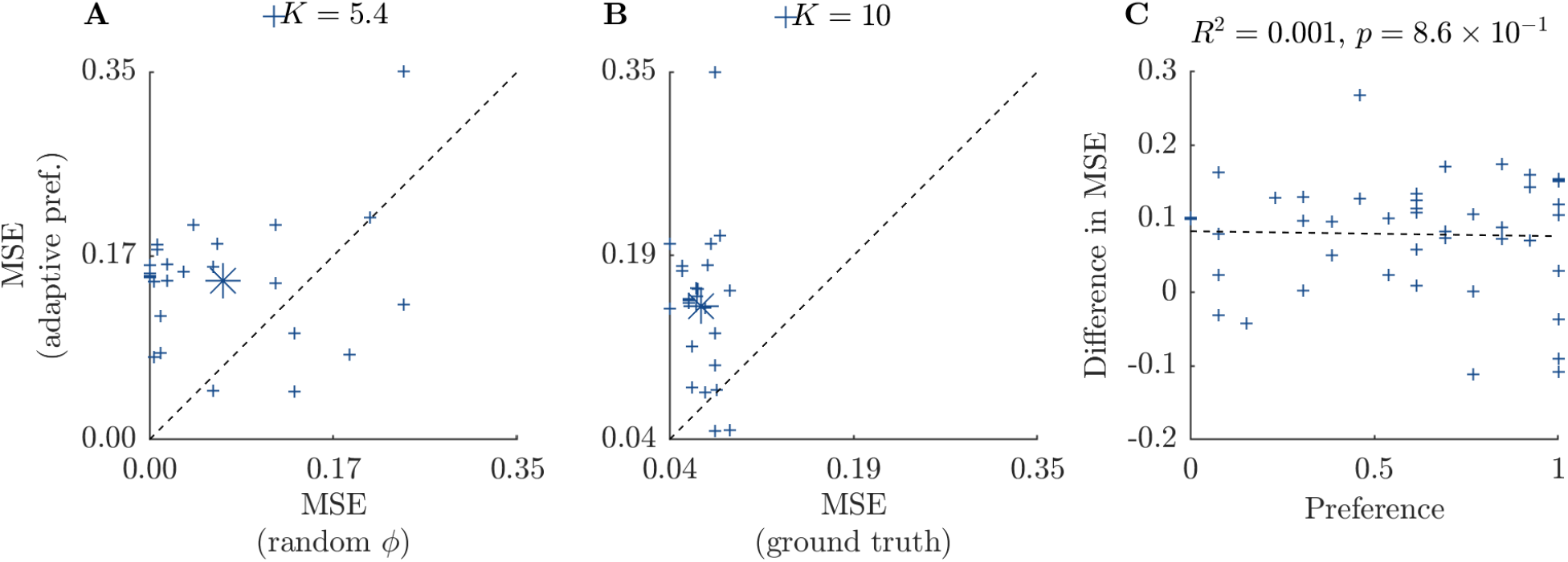
We computed, for each encoder, the MSE between predicted percepts and stimuli for the 26 letters. **A**. The MSE is higher on average for the optimized encoder than for the encoders with random parameters (K = 5.4, decisive evidence). **B**. The MSE is higher on average for the optimized encoder than for the encoders with the true model parameters (K = 10, decisive evidence). The asterisks represent the centroids. **C**. The difference in MSE between two encoders is not correlated with the preference between them (F-test, p = 0.86).

## Notes

### Competing Interest Statement

The authors have declared no competing interest.

## References

A. K. Ahuja, J. D. Dorn, A. Caspi, M. J. McMahon, G. Dagnelie, L. DaCruz, P. Stanga, M. S. Humayun, and R. J. Greenberg. Blind subjects implanted with the Argus II retinal prosthesis are able to improve performance in a spatial-motor task. Br. J. Ophthalmol., 95(4):539–543, 2011.

N. Barnes, A. F. Scott, P. Lieby, M. A. Petoe, C. McCarthy, A. Stacey, L. N. Ayton, N. C. Sinclair, M. N. Shivdasani, N. H. Lovell, H. J. McDermott, and J. G. Walker. Vision function testing for a suprachoroidal retinal prosthesis: Effects of image filtering. J. Neural Eng., 13(3):1–15, 2016.

M. Becker, R. Eckmiller, and R. Huenermann. Psychophysical test of a tunable retina encoder for retina implants. In Int. Jt. Conf. Neural Networks (IJCNN’99), July 10, 1999 - July 16, 1999, volume 1, pages 192–195. IEEE, 1999.

M. Beyeler. Biophysical model of axonal stimulation in epiretinal visual prostheses. arXiv, 2018.

M. Beyeler, G. Boynton, I. Fine, and A. Rokem. pulse2percept: A Python-based simulation framework for bionic vision. In Proc. 16th Python Sci. Conf., pages 81–88. SciPy, 2017a.

M. Beyeler, A. Rokem, G. M. Boynton, and I. Fine. Learning to see again: biological constraints on cortical plasticity and the implications for sight restoration technologies. J. Neural Eng., 14(5):051003, 2017b.

M. Beyeler, D. Nanduri, J. D. Weiland, A. Rokem, G. M. Boynton, I. Fine, G. M. Boynton, and I. Fine. A model of ganglion axon pathways accounts for percepts elicited by retinal implants. bioRxiv, 9(1):1–6, 2019.

E. Bloch and L. da Cruz. The Argus II Retinal Prosthesis System. In Prosthesis. IntechOpen, 2019.

D. H. Brainard. The Psychophysics Toolbox. Spat. Vis., 10(4):433–436, 1997.

E. Brochu, T. Brochu, and N. Freitas. A Bayesian interactive optimization approach to procedural animation design. Comput. Animat. 2010 - ACM SIGGRAPH / Eurographics Symp. Proceedings, SCA 2010, pages 103–112, 2010.

X. Chen, F. Wang, E. Fernandez, and P. R. Roelfsema. Shape perception via a high-channel-count neuroprosthesis in monkey visual cortex. Science (80-.)., 370(6521):1191–1196, 2020.

L. da Cruz, J. D. Dorn, M. S. Humayun, G. Dagnelie, J. Handa, P. O. Barale, J. A. Sahel, P. E. Stanga, F. Hafezi, A. B. Safran, J. Salzmann, A. Santos, D. Birch, R. Spencer, A. V. Cideciyan, E. de Juan, J. L. Duncan, D. Eliott, A. Fawzi, L. C. Olmos de Koo, A. C. Ho, G. Brown, J. Haller, C. Regillo, L. V. Del Priore, A. Arditi, and R. J. Greenberg. Five-Year Safety and Performance Results from the Argus II Retinal Prosthesis System Clinical Trial. Ophthalmology, 123(10):2248–2254, 2016.

J. P. Dmochowski, A. Datta, M. Bikson, Y. Su, and L. C. Parra. Optimized multi-electrode stimulation increases focality and intensity at target. J. Neural Eng., 8(4), 2011.

R. Eckmiller, R. Hünermann, and M. Becker. Exploration of a dialog-based tunable retina encoder for retina implants. Neurocomputing, 26-27:1005–1011, 1999.

R. Eckmiller, D. Neumann, and O. Baruth. Tunable retina encoders for retina implants: why and how. J. Neural Eng., 2(1):S91–S104, 2005.

R. Eckmiller, R. Schatten, and O. Baruth. Portable Biomimetic Retina for Learning, Perception-based Image Acquisition. In 2007 Int. Jt. Conf. Neural Networks, pages 2436–2441. IEEE, 2007.

T. Fauvel and M. Chalk. Efficient Exploration in Binary and Preferential Bayesian Optimization. arXiv, 2021a.

T. Fauvel and M. Chalk. Contextual Bayesian optimization with binary outputs. arXiv, 2021b.

D. Feng and C. McCarthy. Enhancing scene structure in prosthetic vision using iso-disparity contour perturbance maps. Proc. Annu. Int. Conf. IEEE Eng. Med. Biol. Soc. EMBS, pages 5283–5286, 2013.

D. K. Freeman, J. F. Rizzo, and S. I. Fried. Encoding visual information in retinal ganglion cells with prosthetic stimulation. J. Neural Eng., 8(3), 2011.

J. R. Gardner, C. Guo, K. Q. Weinberger, R. Garnett, and R. Grosse. Discovering and exploiting additive structure for Bayesian optimization. Proc. 20th Int. Conf. Artif. Intell. Stat. AISTATS 2017, 54, 2017.

J. R. Golden, C. Erickson-Davis, N. P. Cottaris, N. Parthasarathy, F. Rieke, D. H. Brainard, B. A. Wandell, and E. J. Chichilnisky. Simulation of visual perception and learning with a retinal prosthesis. J. Neural Eng., 16(2):1–23, 2019.

L. E. Grosberg, K. Ganesan, G. A. Goetz, S. S. Madugula, N. Bhaskhar, V. Fan, P. Li, P. Hottowy, W. Dabrowski, A. Sher, A. M. Litke, S. Mitra, and E. J. Chichilnisky. Activation of ganglion cells and axon bundles using epiretinal electrical stimulation. J. Neurophysiol., 118(3):1457–1471, 2017.

T. Guo, C. Y. Yang, D. Tsai, M. Muralidharan, G. J. Suaning, J. W. Morley, S. Dokos, and N. H. Lovell. Closed-loop efficient searching of optimal electrical stimulation parameters for preferential excitation of retinal ganglion cells. Front. Neurosci., 12(MAR):1–12, 2018.

N. Han, S. Srivastava, A. Xu, D. Klein, and M. Beyeler. Deep Learning–Based Scene Simplification for Bionic Vision. arXiv, 1(1), 2021.

A. Horsager, S. H. Greenwald, J. D. Weiland, M. S. Humayun, R. J. Greenberg, M. J. McMahon, G. M. Boynton, and Fine. Predicting Visual Sensitivity in Retinal Prosthesis Patients. Investig. Opthalmology Vis. Sci., 50(4):1483, 2009.

A. Horsager, R. J. Greenberg, and I. Fine. Spatiotemporal interactions in retinal prosthesis subjects. Investig. Ophthalmol. Vis. Sci., 51(2):1223–1233, 2010.

A. Horsager, G. M. Boynton, R. J. Greenberg, and I. Fine. Temporal interactions during paired-electrode stimulation in two retinal prosthesis subjects. Investig. Ophthalmol. Vis. Sci., 52(1):549–557, 2011.

M. S. Humayun, J. D. Dorn, L. da Cruz, G. Dagnelie, J.-A. Sahel, P. E. Stanga, A. V. Cideciyan, J. L. Duncan, D. Eliott, E. Filley, A. C. Ho, A. Santos, A. B. Safran, A. Arditi, L. V. Del Priore, and R. J. Greenberg. Interim Results from the International Trial of Second Sight’s Visual Prosthesis. Ophthalmology, 119(4):779–788, 2012.

N. M. Jansonius, J. Nevalainen, B. Selig, L. M. Zangwill, P. A. Sample, W. M. Budde, J. B. Jonas, W. A. Lagrèze, P. J. Airaksinen, R. Vonthein, L. A. Levin, J. Paetzold, and U. Schiefer. A mathematical description of nerve fiber bundle trajectories and their variability in the human retina. Vision Res., 49(17):2157–2163, 2009.

P. R. Jones. QuestPlus: A MATLAB Implementation of the QUEST+ adaptive Psychometric Method. J. Open Res. Softw., 6, 2018.

R. E. Kass and A. E. Raftery. Bayes Factors. J. Am. Stat. Assoc., 90(430):773, 1995.

M. Kleiner, D. H. Brainard, D. G. Pelli, C. Broussard, T. Wolf, and D. Niehorster. What’s new in Psychtoolbox-3? Perception, 36:S14, 2007.

A. Kupcsik, D. Hsu, and W. S. Lee. Learning Dynamic Robot-to-Human Object Handover from Human Feedback. In Springer Proc. Adv. Robot., volume 2, pages 161–176, 2018.

J. I. Lee and M. Im. Optimal Electric Stimulus Amplitude Improves the Selectivity between Responses of on Versus off Types of Retinal Ganglion Cells. IEEE Trans. Neural Syst. Rehabil. Eng., 27(10):2015–2024, 2019.

C. Li, S. Gupta, S. Rana, V. Nguyen, S. Venkatesh, and A. Shilton. High dimensional Bayesian optimization using dropout. IJCAI Int. Jt. Conf. Artif. Intell., 0:2096–2102, 2017.

H. Lorach, O. Marre, J.-a. Sahel, R. Benosman, and S. Picaud. Neural stimulation for visual rehabilitation: Advances and challenges. J. Physiol., 107(5):421–431, 2013.

H. Lorach, G. Goetz, R. Smith, X. Lei, Y. Mandel, T. Kamins, K. Mathieson, P. Huie, J. Harris, A. Sher, and D. Palanker. Photovoltaic restoration of sight with high visual acuity. Nat. Med., 21(5):476–482, 2015.

R. Lorenz, L. E. Simmons, R. P. Monti, J. L. Arthur, S. Limal, I. Laakso, R. Leech, and I. R. Violante. Efficiently searching through large tACS parameter spaces using closed-loop Bayesian optimization. Brain Stimul., 12(6): 1484–1489, 2019.

Y. H. L. Luo and L. da Cruz. The Argus® II Retinal Prosthesis System. Prog. Retin. Eye Res., 50:89–107, 2016.

L. T. McIntosh, N. Maheswaranathan, A. Nayebi, S. Ganguli, and S. A. Baccus. Deep learning models of the retinal response to natural scenes. Adv. Neural Inf. Process. Syst., (Nips):1369–1377, 2016.

T. P. Minka. A family of algorithms for approximate Bayesian inference. PhD thesis, Massachusetts Institute of Technology, 2001.

M. Mutný and A. Krause. Efficient high dimensional Bayesian optimization with additivity and quadrature fourier features. Adv. Neural Inf. Process. Syst., 2018-Decem(NeurIPS):9005–9016, 2018.

D. Nanduri, I. Fine, A. Horsager, G. M. Boynton, M. S. Humayun, R. J. Greenberg, and J. D. Weiland. Frequency and amplitude modulation have different effects on the percepts elicited by retinal stimulation. Investig. Ophthalmol. Vis. Sci., 53(1):205–214, 2012.

J. B. Nielsen. Systems for Personalization of Hearing Instruments. PhD thesis, Technical University of Denmark, 2015.

S. Nirenberg and C. Pandarinath. Retinal prosthetic strategy with the capacity to restore normal vision. Proc. Natl. Acad. Sci., 109(37):15012–15017, 2012.

D. Palanker, Y. Le Mer, S. Mohand-Said, M. Muqit, and J. A. Sahel. Photovoltaic Restoration of Central Vision in Atrophic Age-Related Macular Degeneration. Ophthalmology, 127(8):1097–1104, 2020.

E. Peli. Testing Vision Is Not Testing For Vision. Transl. Vis. Sci. Technol., 9(13):32, 2020.

D. G. Pelli. The VideoToolbox software for visual psychophysics: Transforming numbers into movies, 1997.

A. Pérez Fornos, J. Sommerhalder, L. da Cruz, J. A. Sahel, S. Mohand-Said, F. Hafezi, and M. Pelizzone. Temporal Properties of Visual Perception on Electrical Stimulation of the Retina. Investig. Opthalmology Vis. Sci., 53(6): 2720, 2012.

V. Rincón Montes, J. Gehlen, S. Ingebrandt, W. Mokwa, P. Walter, F. Müller, and A. Offenhäusser. Development and in vitro validation of flexible intraretinal probes. Sci. Rep., 10(1):1–14, 2020.

J. F. Rizzo. Update on retinal prosthetic research: The Boston retinal implant project. J. Neuro-Ophthalmology, 31 (2):160–168, 2011.

C. P. Robert. Le choix bayésien. Statistique et probabilités appliquées. Springer-Verlag, Paris, 2006.

P. Rolland, J. Scarlett, I. Bogunovic, and V. Cevher. High-dimensional Bayesian optimization via additive models with overlapping groups. Int. Conf. Artif. Intell. Stat. AISTATS 2018, 84:298–307, 2018.

M. Seeger. Notes on Minka’s expectation propagation for Gaussian process classification, 2002.

N. P. Shah and E. J. Chichilnisky. Computational challenges and opportunities for a bi-directional artificial retina. J. Neural Eng., 17(5):055002, 2020.

N. P. Shah, S. Madugula, E. J. Chichilnisky, J. Shlens, Y. Singer, and G. Brain. Learning a neural response metric for retinal prosthesis. bioRxiv, pages 1–13, 2017.

N. P. Shah, S. Madugula, L. Grosberg, G. Mena, P. Tandon, P. Hottowy, A. Sher, A. Litke, S. Mitra, and E. J. Chichilnisky. Optimization of Electrical Stimulation for a High-Fidelity Artificial Retina. In Int. IEEE/EMBS Conf. Neural Eng. NER, volume 2019-March, pages 714–718. IEEE, 2019.

M. J. Spencer, T. Kameneva, D. B. Grayden, H. Meffin, and A. N. Burkitt. Global activity shaping strategies for a retinal implant. J. Neural Eng., 16(2):026008, 2018.

K. Stingl, R. Schippert, K. U. Bartz-Schmidt, D. Besch, C. L. Cottriall, T. L. Edwards, F. Gekeler, U. Greppmaier, K. Kiel, A. Koitschev, L. Kühlewein, R. E. MacLaren, J. D. Ramsden, J. Roider, A. Rothermel, H. Sachs, G. S. Schröder, J. Tode, N. Troelenberg, and E. Zrenner. Interim results of a multicenter trial with the new electronic subretinal implant alpha AMS in 15 patients blind from inherited retinal degenerations. Front. Neurosci., 11(AUG): 445, 2017.

W. R. Thompson. On the Likelihood that One Unknown Probability Exceeds Another in View of the Evidence of Two Samples. Biometrika, 25(3/4):285, 1933.

M. Tucker, E. Novoseller, C. Kann, Y. Sui, Y. Yue, J. Burdick, and A. D. Ames. Preference-Based Learning for Exoskeleton Gait Optimization. arXiv, 2019.

R. Turner, D. Eriksson, M. McCourt, J. Kiili, E. Laaksonen, Z. Xu, and I. Guyon. Bayesian Optimization is Superior to Random Search for Machine Learning Hyperparameter Tuning: Analysis of the Black-Box Optimization Challenge 2020. arXiv, 2021.

P. Twyford, C. Cai, and S. Fried. Differential responses to high-frequency electrical stimulation in on and off retinal ganglion cells. J. Neural Eng., 11(2), 2014.

Z. Wang, C. Li, S. Jegelka, and P. Kohli. Batched high-dimensional Bayesian optimization via structural kernel learning. 34th Int. Conf. Mach. Learn. ICML 2017, 8:5590–5603, 2017.

A. B. Watson. QUEST+: A general multidimensional Bayesian adaptive psychometric method. J. Vis., 17(3):10, 2017.

A. C. Weitz, D. Nanduri, M. R. Behrend, A. Gonzalez-Calle, R. J. Greenberg, M. S. Humayun, R. H. Chow, and J. D. Weiland. Improving the spatial resolution of epiretinal implants by increasing stimulus pulse duration. Sci. Transl. Med., 7(318):1–12, 2015.

M. Zhang, H. Li, and S. Su. High Dimensional Bayesian Optimization via Supervised Dimension Reduction. arXiv, pages 4292–4298, 2019.

Z. Zhao, A. Ahmadi, C. Hoover, L. Grado, N. Peterson, X. Wang, D. Freeman, T. Murray, A. Lamperski, D. Darrow, and T. I. Netoff. Optimization of Spinal Cord Stimulation Using Bayesian Preference Learning and Its Validation. IEEE Trans. Neural Syst. Rehabil. Eng., 29:1987–1997, 2021.

